# Activation and depression of neural and hemodynamic responses induced by the intracortical microstimulation and visual stimulation in the mouse visual cortex

**DOI:** 10.1101/2024.01.01.573814

**Authors:** Naofumi Suematsu, Alberto L Vazquez, Takashi DY Kozai

## Abstract

*Objective*. Intracortical microstimulation can be an effective method for restoring sensory perception in contemporary brain-machine interfaces. However, the mechanisms underlying better control of neuronal responses remain poorly understood, as well as the relationship between neuronal activity and other concomitant phenomena occurring around the stimulation site. *Approach*. Different microstimulation frequencies were investigated *in vivo* on Thy1-GCaMP6s mice using widefield and two-photon imaging to evaluate the evoked excitatory neural responses across multiple spatial scales as well as the induced hemodynamic responses. Specifically, we quantified stimulation-induced neuronal activation and depression in the mouse visual cortex and measured hemodynamic oxyhemoglobin and deoxyhemoglobin signals using mesoscopic-scale widefield imaging. *Main results*. Our calcium imaging findings revealed a preference for lower-frequency stimulation in driving stronger neuronal activation. A depressive response following the neural activation preferred a slightly higher frequency stimulation compared to the activation. Hemodynamic signals exhibited a comparable spatial spread to neural calcium signals. Oxyhemoglobin concentration around the stimulation site remained elevated during the post-activation (depression) period. Somatic and neuropil calcium responses measured by two-photon microscopy showed similar dependence on stimulation parameters, although the magnitudes measured in soma was greater than in neuropil. Furthermore, higher-frequency stimulation induced a more pronounced activation in soma compared to neuropil, while depression was predominantly induced in soma irrespective of stimulation frequencies. *Significance*. These results suggest that the mechanism underlying depression differs from activation, requiring ample oxygen supply, and affecting neurons. Our findings provide a novel understanding of evoked excitatory neuronal activity induced by intracortical microstimulation and offer insights into neuro-devices that utilize both activation and depression phenomena to achieve desired neural responses.

## Introduction

Penetrating microelectrodes have proven invaluable in neurophysiological studies, allowing for the recording [1–4] and stimulation [5–8] of neural activity. They have also shown promising results in neuroengineering applications, particularly in the field of neuroprosthetics [9–11]. For example, in visual cortical prosthetics, intracortical microstimulation (ICMS) delivered through implanted probes in the visual cortex has been extensively used to investigate evoked neural responses [12–14], animal behaviors [15–18], and even restore some aspects of visual perception and perceived visual experiences in human subjects [19–23]. These studies not only provide new insights into the electrophysiological properties of cortical networks but also underscore the need for additional studies to enable successful applications of future visual prosthetic devices. In order to make progress towards these applications, a deeper understanding of the underlying mechanisms that govern the spatiotemporal properties of neuronal activation by different ICMS parameters is crucial. Despite significant advancements [24–30], there are still numerous gaps in our understanding of the stimulation paradigm, as the parameter space associated with ICMS is vast. Many aspects of the interactions between ICMS and local networks also remain largely unexplored.

While many studies have examined the activation of neuronal responses induced by ICMS, fewer studies have characterized the depressive effects of ICMS on the neural response following stimulation [6, 29, 31–33]. Previous research has demonstrated significant depression of neural activity following single or repetitive ICMS, occurring at various time scales ranging from milliseconds to hours. However, due to variations in recording methods and stimulation parameters, there is still a lack of comprehensive understanding of this depression phenomenon. In the context of sensory restoration, such as visual prostheses, it is crucial to consider the critical frequencies of sensory cortical neurons that can enable continuous sensory perception. Exploring whether refractory periods, neuronal depression, adaptation, or active inhibition contribute to the loss of activation in response to ICMS [34] remains an open area for investigation. Developing a better understanding of ICMS-induced depression will facilitate innovations in controlling this effect to ultimately enhance prosthetic performance.

Emerging imaging tools are providing novel insights into ICMS-induced activation and depression, improving traditional technologies like electrophysiology by capturing activity from populations of neurons with spatial information. Concurrent measurements of tissue oxygenation can help determine whether neuronal depression results from metabolic mismatch [35]. Neuronal activation is typically followed by changes in blood supply, altering local oxygen delivery. Intrinsic optical imaging, sensitive to the oxidative state of hemoglobin, provides a convenient method for obtaining these measurements. Additionally, intrinsic imaging aids in distinguishing between bulk fluorescence decreases resulting from increases in hemoglobin absorption and those caused by decreases in calcium ion (Ca^2+^) concentration. Establishing a quantitative relationship between the evoked neuronal response and oxy-/deoxy-hemoglobin hemodynamic signals is essential for estimating oxygen extraction and consumption (metabolic load) vs. stimulation-induced fatigue caused by ICMS [36].

Here, we conducted quantitative analyses of neuronal Ca^2+^ activity and hemodynamic responses to various ICMS frequencies and visual stimulation in Thy1-GCaMP6s mice, which expressed genetically-encoded Ca^2+^ indicator (GCaMP6s) in excitatory neurons [37] using *in vivo* mesoscopic-scale widefield fluorescence and intrinsic imaging. In addition, we also examined somatic and neuropil responses to ICMS *in vivo* with high-resolution two-photon microscopy. We compared the Ca^2+^ and hemodynamic responses between ICMS and visual stimulation conditions. Our findings revealed that visually-evoked Ca^2+^ responses exhibited balanced magnitudes of activation and depression. However, electrically-induced Ca^2+^ responses were dominated by activation, which was frequency-dependent. We observed that hemodynamic responses persisted beyond the period of Ca^2+^ activation extending into the post-stimulus depression phase. Furthermore, our two-photon Ca^2+^ imaging demonstrated that the 25-Hz ICMS resulted in the most pronounced activation and depression, particularly in the soma, but depression was also observed at higher frequencies. These findings indicate that neuronal depression resulting from stimulation-induced reduction in neuronal excitability exhibits a distinct frequency preference compared to activation. This depression effect demands a high energy supply and has a relatively stronger impact on the activated region compared to visual stimulation. Taken together, these results provide novel insights into microstimulation-induced neural activity, which can contribute to the advancement of ICMS strategies for numerous applications including intracortical sensory prostheses.

## Methods

### Animals and surgery

Transgenic mice expressing the Ca^2+^ indicator in excitatory neurons (Thy1-GCaMP6s; 024275, Jackson Laboratory, ME, USA) were used for simultaneous recordings of Ca^2+^ and hemodynamic signals under the mesoscopic-scale microscopy (n = 8) and for two-photon Ca^2+^ imaging (n = 9) studies. Surgical procedures were conducted following previously established protocols [25, 28, 29]. Briefly, mice were anesthetized with a mixture of xylazine (7 mg/kg b.w., i.p.; Covetrus, OH, US) and ketamine hydrochloride (75 mg/kg b.w., i.p.; Covetrus). Craniotomies were then performed over the left visual cortex (AP = –1–-5, LM = 0.5–3.5 mm). Custom stimulation microelectrodes (Q1×4-3mm-50-703-CM16LP, NeuroNexus Technologies) were implanted at an angle of 20–30 degrees slanted towards the posterior region. The electrode was securely fixed to the skull with UV-curable resin (e-on Flowable A2, BencoDental, PA, USA). Reference and ground lines on the electrode connector were wired to the bone screws placed in the skull over the frontal lobes. Kwik-Sil was used to fill the exposed cortical surfaces in the craniotomy to maintain visibility, and a cover glass was used to seal the craniotomy with UV-curable resin. Anesthesia level was periodically monitored and an update (ketamine, 45 mg/kg b.w., i.p.) was administered as necessary. Throughout the surgical procedure and recovery period, the mice’s body temperature was maintained using a heating water pad and respiration was monitored visually. Following surgery, antipamezole hydrochloride (Antisedan, Zoetis, NJ, US; 1 mg/kg b.w., i.p.) and ketoprofen (Ketofen, Zoetis; 5 mg/kg b.w., i.p.) were administered for reversal of anesthesia and for analgesia. After recovery, mice were allowed to recover for up to 2 hours prior to the recording session. Animals were kept in a 12-h/12-h light/dark cycle. All procedures described here were approved by the Division of Laboratory Animal Resources and Institutional Animal Care and Use Committee at the University of Pittsburgh and performed in accordance with the National Institutes of Health Guide for Care and Use of Laboratory Animals.

### Mesoscopic and Microscopic Image Acquisition

Animals were placed in a custom treadmill and head-fixed throughout the imaging experiments. We employed a mesoscopic-scale widefield microscope (MVX-10 epifluorescence microscope; Olympus, Tokyo Japan; 6.3x zoom) for simultaneous recording of Ca^2+^ and intrinsic optical signals over a relatively larger field-of-view of 2.3mm × 1.7mm (Fig. 1a). The imaging was conducted with three sequentially interwoven LED illuminations, and barrier filters were placed in front of each LED light source to restrict the spectral band of each LED to the desired wavelength: 1) blue for GCaMP6s excitation (482.5±15.5 nm; #67-028, Edmund Optics Inc., NJ, US); 2) green for measuring hemoglobin-sensitive imaging (532±2 nm; FLH532-4, Thorlabs Inc., NJ, US); and 3) red for differentiating oxy– and deoxy-hemoglobin (633±2.5 nm; FLH633-5, Thorlabs Inc.) [35, 38]. Fluorescence and reflectance intensities were collected through a long-pass filter (> 500 nm; ET500lp, Chroma Technology Corporation, VT, US) placed in the emission path before the camera (QImaging Retiga R1; Cairn Research Ltd, Kent, UK) and images were sampled at 30 Hz (10 Hz for each color). The camera was controlled using Micro-Manager software [39, 40] with an imaging matrix size of 344 × 256 pixels (Fig. 1b–e).

**Figure 1.**
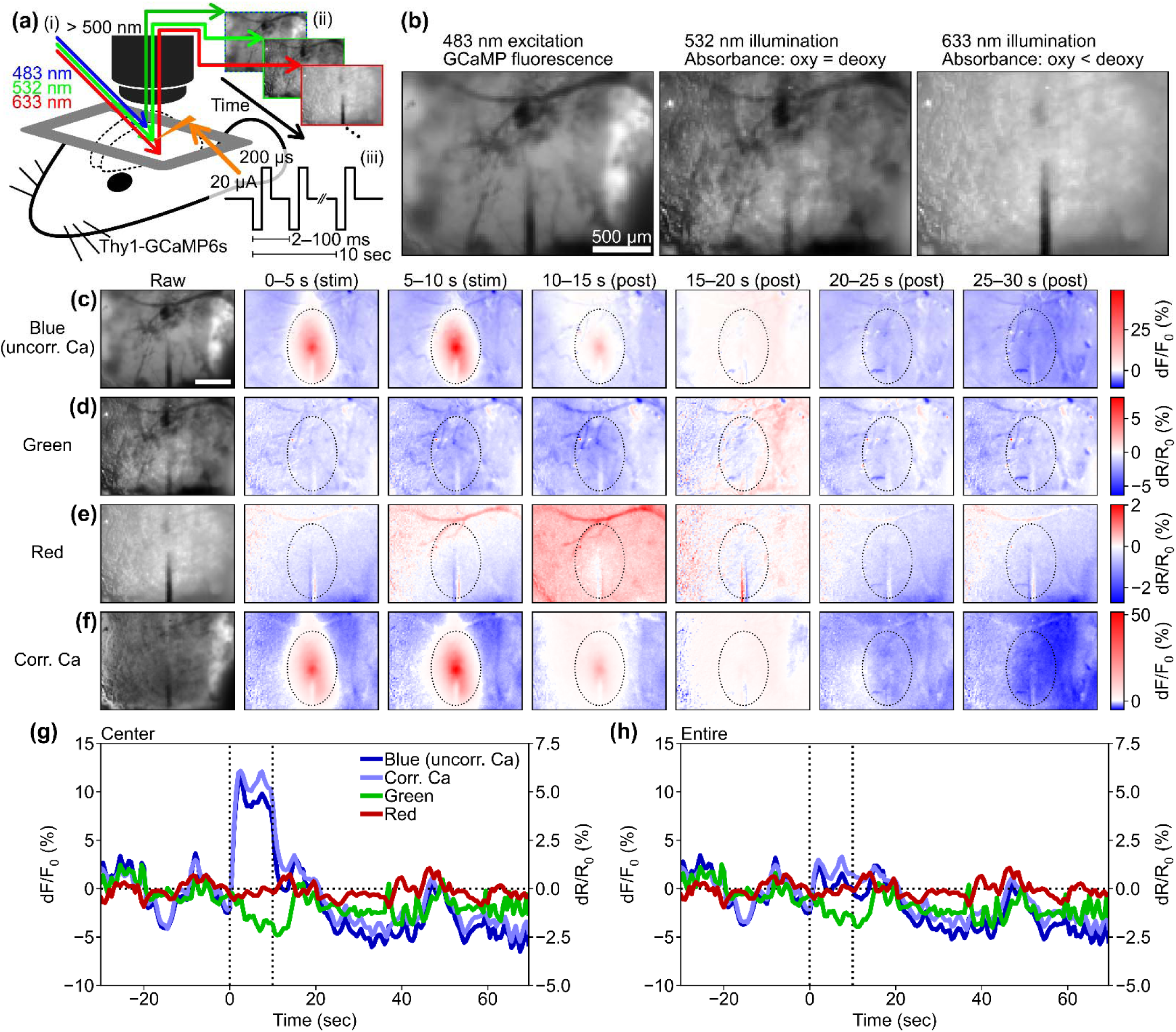
Mesoscopic-scale widefield imaging of Ca^2+^ and hemodynamic signals. (a) The experimental setup Involved applying a widefield single-photon excitation of GCaMP (483 nm) and hemodynamic-visualization illumination (532 and 633 nm) to the mouse cortex (i) and acquiring mesoscopic-scale fluorescent and reflectance images (ii) during ICMS (iii). (b) Example images taken with different colored light sources. GCaMP fluorescence image with 483-nm excitation light (left), total hemoglobin absorbance image with 532-nm illumination (middle), and deoxyhemoglobin-dominant absorbance image with 633-nm illumination (right). (c–f) Raw and change-ratio (0–30 s from the 10-Hz ICMS onset) images with blue (c), green (d), and red (e) illuminations. Blue corresponds to uncorrected Ca^2+^ signals. Vessel patterns are visible in these images. (f) Hemodynamic correction of Ca^2+^ signal attenuates reductions on the vessel patterns. (g) Four different time courses of uncorrected Ca^2+^, corrected Ca^2+^, green, and red signals simultaneously recorded in the 10-Hz ICMS condition within a center region (dotted ellipse in c–f, 400 µm in horizontal and 600 µm in vertical axes from the stimulation site). The hemodynamic correction shifts the Ca^2+^ signals upward during and after the stimulation period. (h) Time courses in the entire imaging field. Scale bars = 500 µm.

Two-photon microscopy (Bruker, Madison, WI, US) was performed using a laser tuned to 920-nm wavelength (Insight DS+, Spectra-Physics, Menlo Park, CA) for GCaMP6s excitation (laser power ≈ 10–20 mW, Figure 11a). Resonant Galvo scanning was performed to capture higher resolution images at 30 Hz over a 413 × 413 µm^2^ field of view (512 × 512 pixels) at a single z-plane focused around the stimulation site. In both imaging conditions, each recording was composed of a 30-s pre-stimulation period, a 10-s stimulation period, and a 60-s post-stimulation period.

### Electrical (ICMS) and Visual Stimulation

ICMS was performed using an IZ2 stimulator controlled by an RZ5D base processor (Tucker-Davis Technologies, FL, US). Each pulse consisted of a 200-µs cathodic leading phase followed by a 200-µs anodic phase, with balanced charges between the cathodic and anodic phases. The current was fixed to ±20 µA (4 nC/phase), selected to be lower than the safety limit (k ≈ 0.36 < 1.7, [41]). Six different train frequencies (10, 25, 50, 100, 250, and 500 Hz) were tested. The order of ICMS trials was repeated two times in ascending-descending frequency order (10 to 500, 500 to 10, 10 to 500, 500 to 10 Hz).

For comparison, a single blue LED was positioned in front of the animal’s eye contralateral to the imaging window as a more natural visual stimulus (n = 5 out of 8 mice in the mesoscopic-scale widefield imaging, n = 5 out of 9 mice in the two-photon imaging). The LED was controlled by the stimulator, applying a square wave at 10 Hz (50-ms on, 50-ms off, 50% duty cycle) with an amplitude of 2.5 V. All stimulation trials were repeated 4 times.

## Analyses and Statistics

### Pre-processing of the widefield and two-photon imaging data

Analyses were performed using custom-made scripts written in Python (ver. 3.9, Python Software Foundation, DE, US) along with existing modules: numpy, scipy, scikit-image, statsmodels, and matplotlib. The frames of image stacks were preprocessed through spatial binning (to 20 µm/pix for the mesoscopic-scale widefield imaging, and to 1 µm/pix for two-photon imaging) and temporal binning (to 2 Hz) with anti-alias filtering to enhance signal-to-noise ratio. Image registration was conducted on each frame to align them to the first frame of each acquisition by cross-correlation with sub-pixel estimation. For the two-photon microscopy images, additional inter-line alignment was applied as necessary to correct for shearing between lines.

For the widefield imaging data, masks were generated to exclude blood vessels from further analyses (Fig. S1). Average images from each color (blue, green, and red) were intensity-threshold binarized using hysteresis thresholding with the following two threshold levels: (*i*) mean-0.5 standard deviation (SD) and (*ii*) mean-1SD. Spatially connected vessel patterns were extracted after the hysteresis thresholding. A pixel-wise logical summation of all color channels was then calculated to combine them and generate the final vessel pattern image. This procedure also captured pixels on probe shanks that exhibited noticeably low fluorescence and reflectance values.

### Somatic region-of-interest (ROI) definition for the two-photon imaging data

Response images of excitatory neurons acquired under the two-photon microscopy were constructed by calculating pixel-wise mean fluorescent change ratio during the stimulation period (0–10 s) over pre-stimulation mean fluorescent intensity across the entire imaging region. For each animal, these response images for all stimulation conditions were averaged. Region of interest (ROI) for each soma was manually defined with Fiji [42].

### Neuronal Ca^2+^ response quantification for widefield and two-photon imaging

To quantify the excitatory Ca^2+^ activity for two-photon imaging, the fluorescence change ratio (dF/F_0_ = (F(t) – F_0_)/F_0_) was calculated, where F(t) represents the fluorescent intensity at time t and F_0_ is the mean fluorescent intensity over the pre-stimulation period (–30 ≤ t < 0, in s).

In the case of the mesoscopic-scale widefield imaging, the Ca^2+^ signals were corrected based on the green channel values to account for the hemodynamic effect that could potentially affect the green GCaMP signals [35, 38] (Fig. 1f–h). The corrected dF/F_0_(t) was calculated using the formula:

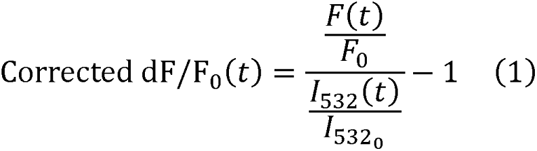

where *I*_532_ (t) represents the reflectance intensity to the green illumination at a specific time point *t*, and *I*_5320_ is the mean of the reflectance intensity to the green illumination during the pre-stimulation period.

### Hemodynamic activity quantification of mesoscopic-scale widefield imaging

The change in oxy– and deoxy-hemoglobin concentrations relative to baseline (and, respectively) were calculated as follows [35]:

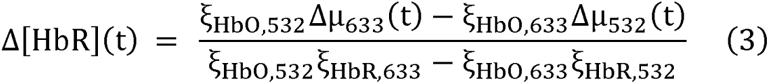

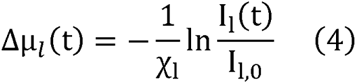

where constant ξ represents the absolute absorption spectra of HbR and HbO at 532 and 633 nm, constant χ represents the pathlengths at wavelength l (i.e., l = 532 and 633 nm). I_l_(t) and I_l,0_ are the reflectance intensities at a time point t and its mean value during the pre-stimulation period for wavelength l. The values of ξ and χ were obtained from [35], and if an exact wavelength value was not provided, the mean value of the adjacent two values was used.

The change in the total hemoglobin concentration (Δ[HbT](t)) was calculated by summing the changes in oxy– and deoxy-hemoglobin (Δ[HbT](t) == Δ[HbO](t) + Δ[HbR](t)). The oxygen extraction fraction (OEF) was calculated using the following:

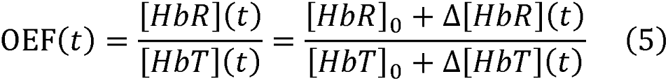

where [HbR]_0_ and [HbT]_0_ represent baseline concentrations of the deoxyhemoglobin and total hemoglobin. For our calculations, we assumed baseline concentrations of 2 mM for hemoglobin in blood, 3% of blood in brain tissue (60 µM of total hemoglobin), and 75% of oxygenation (15 µM of deoxyhemoglobin) [35]. OEF values range between 0 and 1, and values outside this range were excluded as outliers.

### Extraction of significant activation and depression in the widefield imaging data

To define activation and depression distances (from stimulation site), latencies (onset timing), and durations (offset timing – onset timing) of the spatiotemporal magnitude profile of the corrected dF/F_0_, Δ[HbO], and Δ[HbR] while reducing the effect of data fluctuations, we employed hysteresis thresholding. As described above, this method requires two different threshold levels [43]. First, the original three-dimensional data (dF/F_0_, Δ[HbO], and Δ[HbR]) obtained under the mesoscopic-scale imaging (X × Y × time) was converted into two-dimensional structure (distance from stimulation site × time). We calculated the mean ± 1 and 2 SDs of the pre-stimulation period (–30–0 s), and used them as the soft and hard thresholds. To determine the activation distance, we identified points on the distance profile with values above 2 SDs as well as points between 1 and 2 SDs that were connected to distances above 2 SDs as above threshold. The same procedure was applied in the time domain, resulting in a joint threshold image (distance × time) that contained multiple continuous regions. The regions extracted by the hysteresis thresholding were considered candidates for significant activation and depression. We selected at most one region for each activation and depression event based on specific criteria. For activation candidates in ICMS experiments, we selected single regions that (1) emerged within 10 s after stimulation onset, (2) occurred within 50 µm from the stimulation site, and (3) had the longest duration among the candidates. For depression candidates in ICMS conditions, we selected single regions that (1) emerged within 20 s after stimulation onset, (2) occurred within 50 µm from the stimulation site, and (3) had the longest duration among the candidates. Similarly, for visual stimulation conditions, we used criteria (1) and (3) regardless of whether the responses originated from the inserted probe. The onset timing of the extracted region is defined as latency, and the difference between onset and offset timings is duration. Activation and depression distances and mean amplitude across the distance are represented as functions of time from stimulation onset.

After defining the distance, latency, and duration of significant Ca^2+^ activation or depression and hemodynamic activities (d[HbO] and d[HbR]) using the hysteresis thresholding method described above, these measurements were averaged across time for each mouse (e.g., symbols in Figure 4d). Also, the averaged measurements for each mouse were averaged across mice to evaluate these effects as a function of stimulation conditions such as ICMS frequencies and visual stimulation (e.g., bars in Figure 6).

The balance of activation and depression was quantified using the following metrics: distance from the stimulated site, response magnitude, and response duration. Balance based on these metrics was calculated as follows:

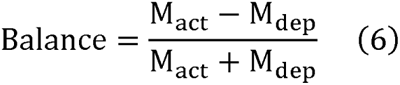

where M_act_ and M_dep_ represent metrics for activation and depression, respectively. The resulting score ranged from –1 (biased to depression) to 1 (biased to activation); 0 indicated balanced activation and depression.

### Statistical procedures

For pair-wise comparisons, we conducted t-tests on two-related samples (e.g., Figure 3c). One-way and two-way analysis of variance (ANOVA) were used for a comparison among three (or more) groups. For example, in Figure 5a, we compared neural activation distances across different stimulation conditions using one-way ANOVA, and in Figure 9a, we compared response distances across stimulation conditions and signal modalities using two-way ANOVA. Post-hoc analysis was performed using Tukey’s honestly significant difference (HSD) test. To quantify correlations between signals from shared units (e.g., between dF/F_0_ values in % or |%| in Figure 5h), the Pearson correlation coefficient was used. For correlations involving signals with different units (e.g., between dF/F_0_ in % and oxyhemoglobin concentration in µM in Figure S4e), we used Spearman’s rank correlation coefficient. Statistical significance was evaluated based on a significance level of *p* < 0.05. Significant results are denoted by asterisk symbols in the figures or summarized in tables. In cases where there were samples without significant activation and/or depression, the metrics such as magnitude and duration were replaced with zero in the statistical analyses of the two-photon imaging data.

## Results

In this study, we investigated the spatiotemporal dynamics of excitatory neurons, focusing on their activation and depression, using 10-s ICMS trains at various frequencies and compared those to visual stimulation. These experiments were conducted using widefield imaging and two-photon microscopy. We quantified the magnitude, spatial extent, latency, and duration of the excitatory neuron responses as well as the induced oxyhemoglobin and deoxyhemoglobin signals. Throughout this paper, we refer to “depression” as the phenomenon in which the neural Ca^2+^ signal decreases in intensity due to stimulation compared to the pre-stimulus period.

### ICMS elevates excitatory Ca^2+^ activity followed by post-stimulus depression

Although stimulation-induced depression of neuronal excitability has been previously described [32], most imaging studies have focused on stimulation-induced activation [5, 24, 27, 44]. To investigate the extent of spatiotemporal activation and depression, we first analyzed ICMS-induced changes in Ca^2+^ activity within excitatory neurons using widefield imaging. We first show results from a representative animal to highlight the spatio-temporal changes (Fig. 2; see Movies S1–S3 for entire frames). A 10-s long ICMS train at 10 Hz was delivered to the visual cortex and produced a clear increase in Ca^2+^ activity during the stimulation period, which spread to approximately 500 µm away from the stimulation site (0–10 s; Figure 2a–c). Following the cessation of the stimulation train (at 10 s), the Ca^2+^ activity rapidly declined and dipped below pre-stimulus baseline (“depression”) approximately 5 s after the ICMS offset (Fig. 2b) and persisted for several tens of seconds during the post-stimulation period although with a noticeably smaller magnitude than activation. This depression effect was particularly prominent in the region exhibiting an increase in Ca^2+^ (Fig. 2c). The Ca^2+^ data were corrected for hemodynamic absorption (as shown in Figure 1f–h) to mitigate this potential signal source. Similar patterns of activation and depression were also observed with 100-Hz ICMS (Fig. 2d–f), but no such effects were observed during visual stimulation (Fig. 2g–i), where the visual stimulation train induced transient activation at the onset and offset periods. Visually-evoked activation and depression spread through almost the entire imaging field due to the use of uniform visual field stimulation.

**Figure 2.**
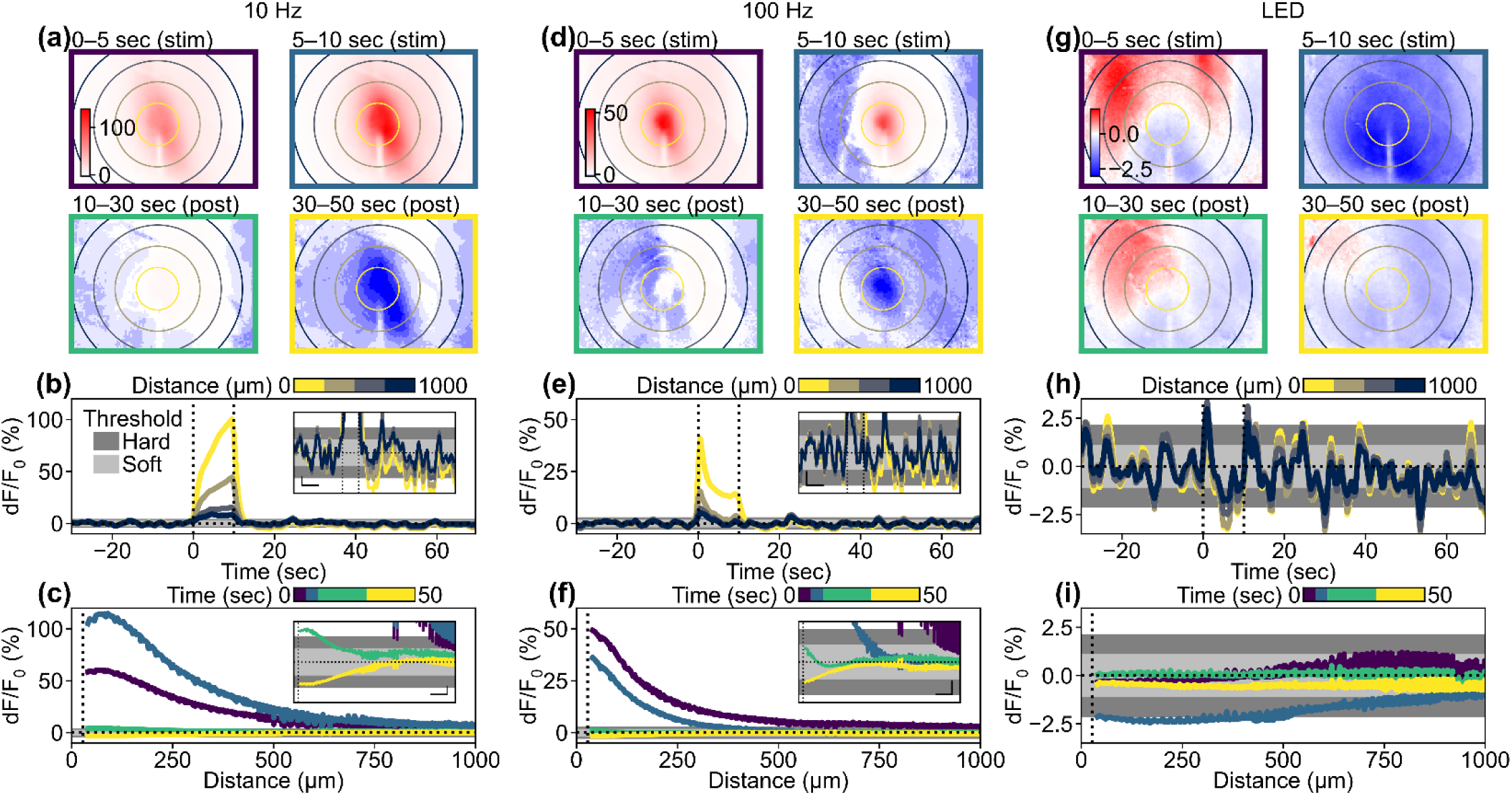
Activation followed by depression of Ca^2+^ responses induced by stimulation. (a, d, and g) Single-animal examples of colormap in the X-Y domain during (a) 10-Hz ICMS, (d) 100-Hz ICMS, and (g) visual stimulation. Concentric circles indicate distances from the stimulation site, plotted at 250-µm intervals. Asymmetric color scales were used for enhanced visualization. (b, e, and h) Time courses of the Ca^2+^ responses to (b) 10-Hz ICMS, (e) 100-Hz ICMS, and (h) visual stimulation at various distances from the stimulation site. Line colors correspond to ones of concentric circles in (a), (d), and (g). Light and dark gray areas indicate pre-stimulation fluctuations (mean ± 1SD and mean ± 2SD) used for the hysteresis thresholding. Vertical dotted lines indicate stimulation onset (t = 0 s) and offset (t = 10 s). Horizontal dotted lines indicate pre-stimulation baseline (d /F_0_ = 0). Insets indicate enlarged time courses (scalebars for x and y axes = 10 s and 1%, respectively). (c, f, and i) Temporal changes in distance profiles of Ca^2+^ response to (c) 10-Hz ICMS, (f) 100-Hz ICMS, and (i) visual stimulation. Line colors correspond to ones of panel frames in (a), (d), and (g). Vertical dotted lines indicate half of the probe width (27.5 µm), where data points are excluded due to vessel masking. Insets indicate enlarged distance profiles (scalebars for x and y axes = 100 µm and 1%, respectively).

The magnitude, duration, and distance of activation and depression were quantified for each animal. The Ca^2+^ responses induced by the 10-Hz ICMS across all animals (Fig. 3a–c) showed that the average magnitude of Ca^2+^ activation increased over time, while the average magnitude of Ca^2+^ depression remained relatively low and constant (Fig. 3a). The activation distance remained constant throughout the ICMS train, followed by varying degrees of depression distance ranging from 300–1000 µm away from the stimulation sites (Fig. 3b). Significant activation was observed in all mice (8/8), while significant depression was observed in most cases (5/8). Although the depression magnitude was attenuated by the hemodynamic correction (paired t-test, *t* = 1.75, *p* = 0.12), it was significantly greater than zero (one-sample t-test, *t* = 3.03, *p* < 0.05; Figure S2a). The depression component in the uncorrected Ca^2+^ signals was significantly stronger than the change obse ved in the green channel (paired t-test *t* = 3.00, *p* < 0.05; Figure S2b). These observations show that the hemodynamic correction works sufficiently with our data and the Ca^2+^ depression remained even after the hemodynamic correction, meaning that the depression was not a feint due to hemodynamic signal contamination.

**Figure 3.**
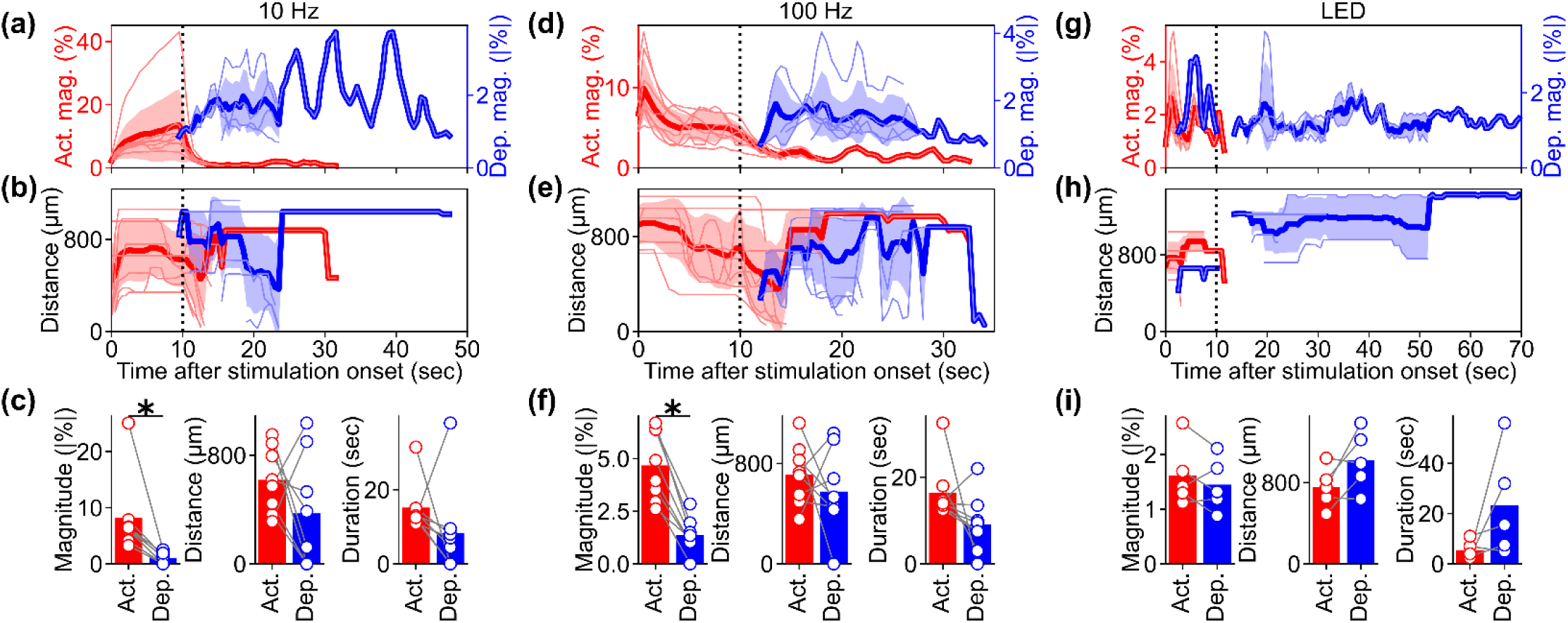
Population analyses of Ca^2+^ activation and depression induced by stimulation. (a) Temporal dynamics of Ca^2+^ activation (red, left vertical axis) and depression (blue, right vertical axis) magnitudes under 10 Hz ICMS condition. Each pale line indicates an individual mouse. Thick lines and shaded areas indicate mean ± 1SD. (b) Temporal dynamics of the activation (red) and depression (blue) distances under 10-Hz ICMS condition. (c, left) Comparison of mean magnitudes be ween the activation and depression under 10-Hz ICMS condition. (c, middle) Comparison of mean distances between activation and depression under 10-Hz ICMS condition. (c, right) Comparison of durations between the activation and depression under 10-Hz ICMS condition. (d–f) Same metrics evaluated under 100-Hz ICMS condition. (g–i) Same metrics evaluated under visual stimulation condition.

The Ca^2+^ activation was significantly stronger than the depression by almost an order of magnitude (comparison of absolute magnitude; Figure 3c, left; *t* = 3.22, *p* < 0.05). However, there was no statistical difference in the mean distances between activation and depression (Fig. 3c, middle; *t* = 1.29, *p* = 0.24). The duration of the activation tended to be longer than the duration of depression (Fig. 3c, right; *t* = 1.22, *p* = 0.26). It is worth noting that the Ca^2+^ activation lasted slightly longer than the 10-s stimulation period. These results indicate that the 10-s, 10-Hz ICMS train consistently evoked sustained activation throughout the stimulation period, followed by depression to a similar spatial extent. These trends were highly dependent on the stimulation parameters and modalities, where 100-Hz ICMS induced transient activation that was stronger than depression in magnitude (Fig. 3d–f, *t* = 7.97, *p* < 0.05), and visually-induced activation was comparable to depression (Fig. 3g–i, *t* = 1.07, *p* = 0.34).

### ICMS frequencies modulate Ca^2+^ activation and depression

Given that ICMS-induced activation is followed by depression of the Ca^2+^ response, we then asked how the stimulation frequencies can impact features of the induced activation and depression of the Ca^2+^ response (Fig. 4). In a representative animal, we observed activation-dominant responses to the ICMS where lower-frequency stimulation induced an elevating (or sustained) response and higher-frequency stimulation elicited a transient and localized response. Meanwhile, the visually-evoked response exhibited transient activation and depression at similar magnitudes. These representative examples of the excitatory neural Ca^2+^ responses provide valuable inspiration about the complex interactions between stimulation frequencies and their effects on the Ca^2+^ response.

**Figure 4.**
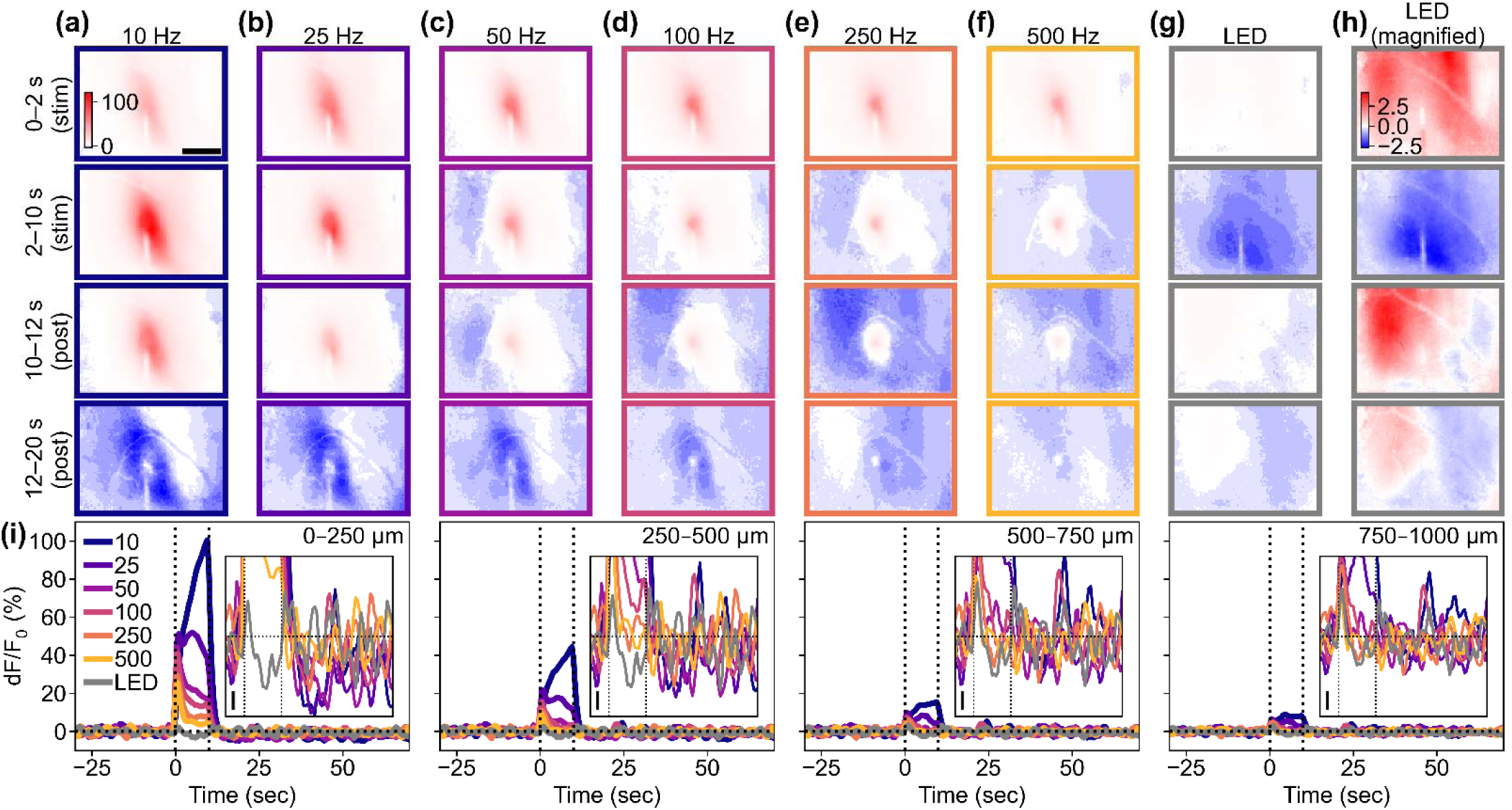
Frequency dependency of ICMS-Induced responses and comparison with visual-evoked response. (a–f) Responses to ICMS with various frequencies. (First and second rows): Transient (first row) and sustained (second row) phases during the stimulation period. (Third and fourth rows): Transient (third row) and sustained (fourth row) phase after the stimulation train offset. Scale bar = 500 µm. (g) Response to 10-Hz LED blink stimulation to the contralateral eye. The color scales in (a) to (g) are the same for comparison. (h) Response to the visual stimulation shown with a magnified color scale for better visibility. (i) Time courses of the ICMS-induced and visual responses at different distances from the stimulation site (left to right: 0–250, 250–500, 500–750, and 750–1000 µm). Insets show the responses around the onset and offset of the stimulation train (vertical dotted lines, 10 s) with a magnified y scale (scale bar = 1% dF/F_0_; horizontal dotted line = 0%).

### Ca^2+^ activation and depression prefer different ICMS frequencies

Given the observed frequency-dependent relationship between the magnitude, area, and spatiotemporal aspects of ICMS-induced activation and depression, we conducted further analysis to quantify the differences between stimulation conditions and examine the correlations between activation and depression (Fig. 5). We determined the response distances and durations of significant activation and depression using the hysteresis thresholding of the mesoscopic-scale Ca^2+^ spatiotemporal dynamics (see Methods, “Extraction of significant activation and depression in the mesoscopic-scale imaging data”). Neither the activation and depression distances did not show any statistical differences among the stimulation conditions (Fig. 5a, activation, one-way ANOVA, *F* = 1.71, *p* = 0.14; Figure 5b, depression, *F* = 1.50, *p* = 0.20). The balance of the activation and the depression distances (Equation 6) did not differ among the stimulation conditions (Fig. 5c, one-way ANOVA, *F* = 0.65, *p* = 0.69). Furthermore, we did not observe any significant correlations between the activation and the depression distances (Fig. 5d).

Although the area of ICMS-induced activation did not differ significantly, the magnitudes of activation were significantly different among the stimulation conditions (Fig. 5e, one-way ANOVA, *F* = 3.10, *p* < 0.05). Lower-frequency ICMS tended to induce stronger activation, while the visual stimulation tended to elicit weaker activation compared to ICMS (post-hoc Tukey’s HSD test, 10-Hz ICMS vs visual stimulation, difference = – 6.6%, *p* < 0.05). In contrast, the magnitudes of depression were not statistically different across all stimulation conditions (Fig. 5f, one-way ANOVA, *F* = 1.39, *p* = 0.24). Consequently, the balance between these two magnitudes were significantly different among the stimulation conditions (Fig. 5g, one-way ANOVA, *F* = 7.02, *p* < 0.05), with visual stimulation showing balanced activation and depression responses (post-hoc Tukey’s HSD test, 10-Hz ICMS vs visual stimulation, difference = –0.75, *p* < 0.05; 25-Hz ICMS vs visual stimulation, difference = –0.52, *p* < 0.05; 50-Hz ICMS vs visual stimulation, difference = –0.56, *p* < 0.05; 100-Hz ICMS vs visual stimulation, difference = –0.52, *p* < 0.05; 250-Hz ICMS vs visual stimulation, difference = –0.61, *p* < 0.05; 500-Hz ICMS vs visual stimulation, difference = –0.67, *p* < 0.05). Moreover, the magnitudes of activation and depression exhibited significantly strong positive correlation in the 10-, 50-, and 100-Hz ICMS conditions (Fig. 5h; Pearson correlation coefficient, 10-Hz ICMS, *r* = 0.73, *p* < 0.05; 50-Hz ICMS, *r* = 0.82, *p* < 0.05; 100-Hz ICMS, *r* = 0.75, *p* < 0.05). These findings indicate that ICMS at lower frequencies elicits stronger Ca^2+^ activation in excitatory neurons, while visual stimulation induces a balanced response of activation and depression. Additionally, the correlations of magnitudes between activation and depression suggest that there may be shared or related mechanisms underlying the processes of activation and depression in certain ICMS conditions.

**Figure 5.**
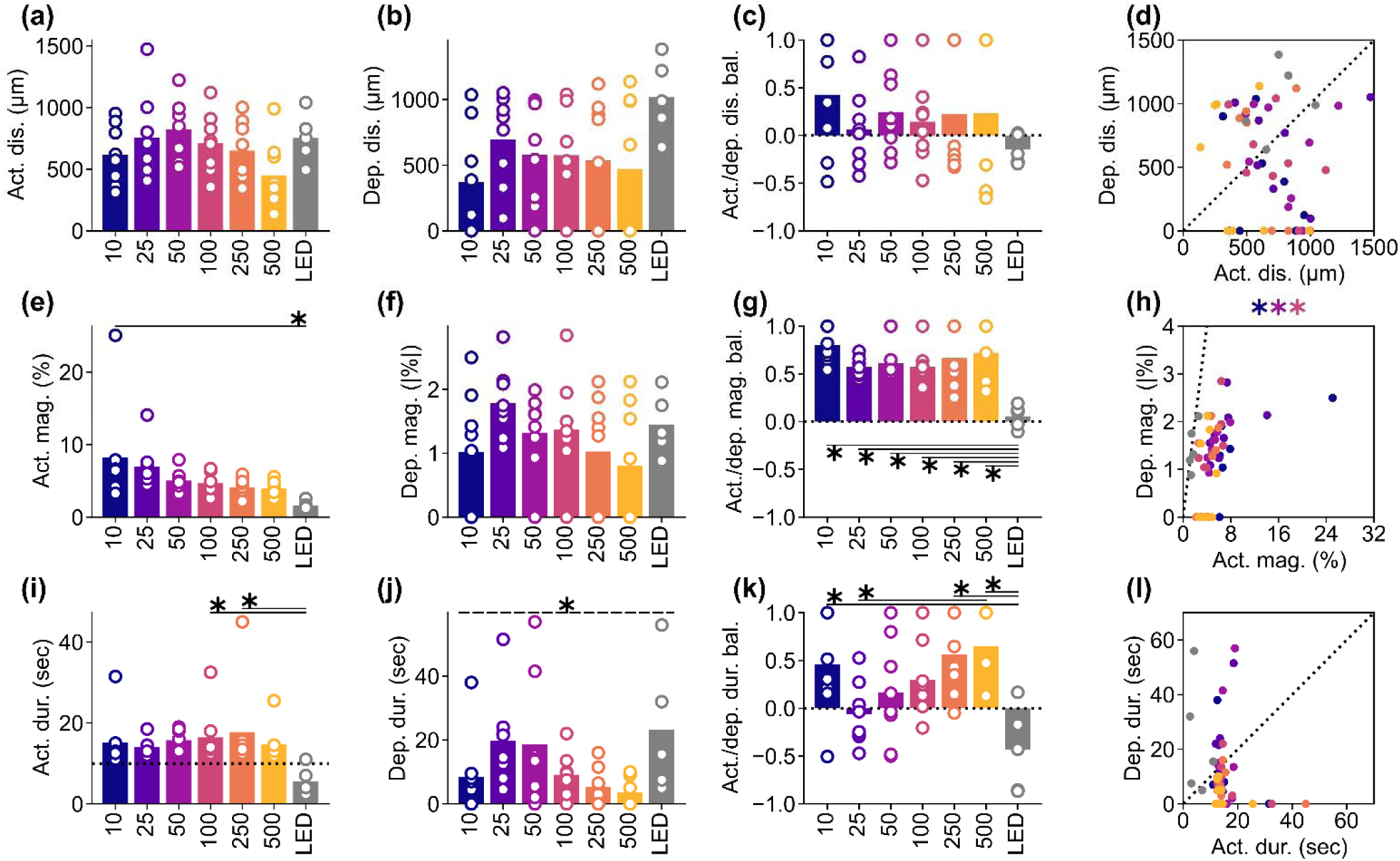
Frequency dependency of the activation and depression distance, magnitude, and duration among various ICMS frequencies and visual stimulation. (a and b) Activation (a) and depression (b) distances. For the ICMSs, the longest-lasting Ca^2+^ response that occurred within 10 s (for activation) and 20 s (for depression) after the stimulation onset, within 50 µm from the stimulation site was extracted. For the visual stimulation, the criterion of stimulation site coverage was not applied. (c) Balance between activation and depression distance (Equation 6). (d) Correlation between activation and depression distances for each animal. (e and f) Activation (e) and depression (f) magnitudes. (g) Balance between activation and depression magnitudes. (h) Correlation between activation and depression magnitudes. (i and j) Activation (i; horizontal dotted line = 10-s stimulation duration) and depression (j) durations. (k) Balance between activation and depression durations. (l) Correlation between activation and depression durations. * with solid line indicates *p* < 0.05 for pairwise comparison using post-hoc Tukey’s HSD test (a–c, e–g, and i–k), * with dashed line indicates *p* < 0.05 for one-way ANOVA without post-hoc pairwise significant difference. Colored * indicates *p* < 0.05 for Pearson correlation coefficient in corresponding conditions (d, h, and l).

The activation durations were significantly dependent on the stimulation conditions (Fig. 5i, one-way ANOVA, *F* = 2.36, *p* < 0.05). All ICMS frequencies, particularly around 100–250 Hz, caused prolonged activation that lasted beyond the duration of the stimulation train (10 s, horizontal dotted line in Figure 5i). In contrast, visual stimulation only caused transient activation at the onset and offset of the stimulus train (post-hoc Tukey’s HSD test, 100-Hz ICMS vs visual stimulation, difference = –11 s, *p* < 0.05; 250-Hz ICMS vs visual stimulation, difference = –12 s, *p* < 0.05). Similar to activation, the duration of depression varied depending on the stimulation conditions (Fig. 5j, one-way ANOVA, *F* = 2.48, *p* < 0.05), with tendencies of longer depression at 25–50-Hz ICMS and visual stimulation compared to the other conditions (no significant pairwise difference by post-hoc Tukey’s HSD test). Consequently, the balance between the activation and depression durations exhibited significant dependence on the stimulation conditions (Fig. 5k, one-way ANOVA, *F* = 4.77, *p* < 0.05). Depression tended to be longer in the 25-Hz ICMS and visual stimulation compared to the other conditions, while activation dominated in the lower and higher-frequency ICMS conditions, especially at 500 Hz (post-hoc Tukey’s HSD test, 10-Hz ICMS vs visual stimulation, difference = –0.89, *p* < 0.05; 25-Hz ICMS vs 500-Hz ICMS, difference = 0.71, *p* < 0.05; 250-Hz ICMS vs visual stimulation, difference = –1.00, *p* < 0.05; 500-Hz ICMS vs visual stimulation, difference = –1.08, *p* < 0.05). No significant correlation was found between the duration of activation and depression (Fig. 5l). These results suggest that ICMS at any frequency entrains the Ca^2+^ activity, while the degree of subsequent depression is dependent on the stimulation conditions. Together, these quantitative findings inform the selection of stimulation frequencies, where the neural responses include both activation and depression phenomena.

### Different ICMS frequencies evoke different spatiotemporal dynamics of Ca^2+^ activation

Since observed that lower-frequency ICMS caused greater sustained Ca^2+^ activation, while higher-frequency ICMS caused transient and localized activation followed by weak sustained activation (Fig. 4i), we proceeded to investigate how activation developed during the stimulation train across different conditions (Fig. 6). In the 10-Hz ICMS condition, the activation gradually increased in magnitude over the 10-s ICMS duration and spread through the visual cortex. On the other hand, in the 100-Hz condition, the Ca^2+^ activity initially peaked, and then decreased over the 10-s ICMS duration, becoming more localized (Fig. 6a–d).

**Figure 6.**
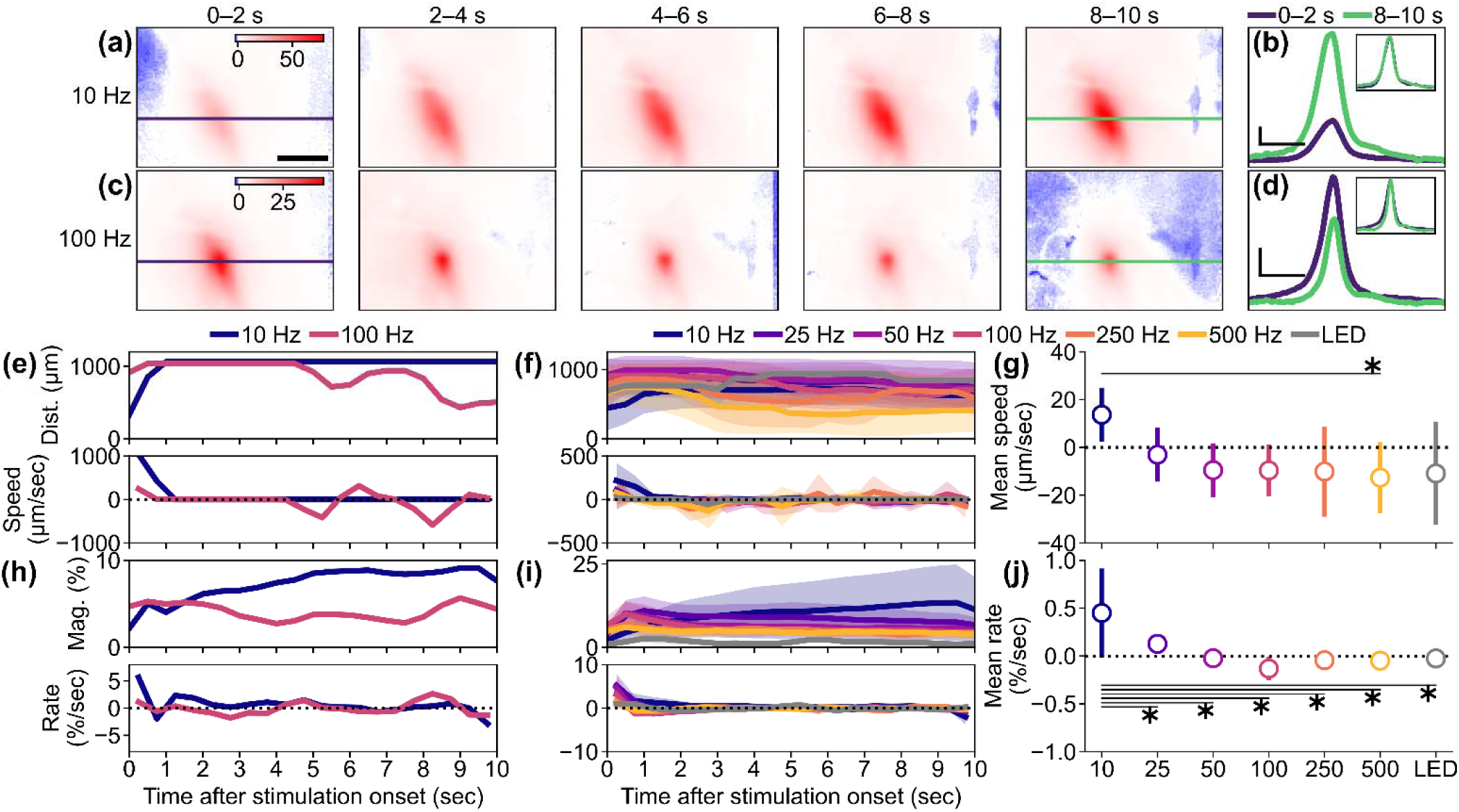
Temporal dynamics of the activations. (a and b) Response maps (a) at 0–2, 2–4, 4–6, 6–8, and 8–10 s after the stimulation onset and line profiles in the medio-lateral axis around the stimulation site (b) at 0–2 (blue) and 8–10 s (green) in response to the 10-Hz ICMS. Scale bar in (a) = 500 µm. Horizontal and vertical scale bars in (b) = 500 µm and 10%, respectively. The Inset in (b) shows the same profiles normalized to each peak value. (c and d) Response maps (c) and line profiles (d) in response to the 100-Hz ICMS. (e) Examples of activation distance (top) a d speed (bottom) during the stimulation period in the 10– and 100-Hz ICMS conditions. Positive and negative speeds indicate expansion and shrinkage of activation distance, respectively. (f) Population activation distance (top) and speed (bottom) (mean ± 1SD). (g) Speed averaged across animals (mean ± 1SD). (h) Examples of the activation magnitude (top) and change rate (bottom). A positive change rate indicates growing-up activation magnitude. A change rate of zero indicates constant activation magnitude. (i) Population magnitude (top) and change rate (bottom) (mean ± 1SD). (j) Magnitude change rate averaged across animals (mean ± 1SD). * with line indicates *p* < 0.05 for pairwise comparison with post-hoc Tukey’s HSD test (g and j).

To quantify the spatiotemporal dynamics of neural Ca^2+^ activation, we measured the distance and calculated the spread/withdrawal speed based on the inter-frame difference of the distance (0.5-s interval). For the 10-Hz condition, the activation distance remained relatively stable over the 10-s ICMS period, resulting in a near-zero spreading speed (Fig. 6e). In contrast, the activation distance rapidly decreased after 2 s of the 100-Hz ICMS, leading to a negative speed (withdrawal of activation) observed 4–5 s after the onset. The mean spread/withdrawal speeds significantly varied across the stimulation conditions (Fig. 6f and g, *F* = 2.78, *p* < 0.05). Specifically, the speed tended to be positive in the 10-Hz ICMS, while it was negative in the other stimulation conditions (post-hoc Tukey’s HSD test, 10-Hz ICMS vs 500-Hz ICMS, difference = –26 µm/sec, *p* < 0.05). This indicates that, apart from the 10-Hz ICMS frequency, the Ca^2+^ activation area gradually withdrew after the initial transient spread in most stimulation conditions.

We next investigated whether the activation magnitude followed the same trend. Analysis of the magnitudes revealed that the 10-Hz ICMS induced an increase in magnitude over the stimulation period, while the 100-Hz ICMS condition showed a relatively constant magnitude (Fig. 6h, top). The change rates of magnitude between each time point remained mostly positive (indicating increasing magnitude) in the 10-Hz conditions, whereas the change rate fluctuated around zero (indicating stable magnitude) in the 100-Hz ICMS condition (Fig. 6h, bottom). The mean change rates of magnitude showed significant differences among the stimulation conditions (Fig. 6i and j, one-way ANOVA, *F* = 6.99, *p* < 0.05). Specifically, lower-frequency ICMS induced elevating activation magnitude (post-hoc Tukey’s HSD test, 10-Hz ICMS vs 25-Hz ICMS, difference = –0.3 %/sec, *p* < 0.05; 10-Hz ICMS vs 50-Hz ICMS, difference = –0.5 %/sec, *p* < 0.05; 10-Hz ICMS vs 100-Hz ICMS, difference = –0.6 %/sec, *p* < 0.05; 10-Hz ICMS vs 250-Hz ICMS, difference = –0.5 %/sec, *p* < 0.05; 10-Hz ICMS vs 500-Hz ICMS, difference = –0.5 %/sec, *p* < 0.05; 10-Hz ICMS vs visual stimulation, difference = –0.5 %/sec, *p* < 0.05). Also, individual neurons exhibited similar temporal dynamics and dependency on stimulation conditions (Fig. S5). Taken together, our findings demonstrate that the lower-frequency ICMS increases both activation area and magnitude over the stimulation period, whereas the higher-frequency ICMS causes a decrease in the activation area and a relatively constant magnitude.

### Ca^2+^ depression emerges distally and then spreads proximally toward stimulation site

Given the stimulation-parameter-dependent spatiotemporal dynamics of activation during sustained ICMS, we next investigated the spatiotemporal dynamics of the depression (Fig. 7). As shown in previous analyses (e.g., Figure 4a–f), the depression onset was initiated in a distal location from the stimulation site. To quantify the spatiotemporal dynamics of depression, we defined the nearest ‘edge distance’ as the distance to the closer edge of the depression to stimulation site by conducting the hysteresis thresholding (see Methods). When examining the nearest edge distances, all ICMS conditions exhibited a decrease in the first few seconds of depression, especially at higher ICMS frequencies (Fig. 7a). This indicated that the depression initially occurred in distal regions and then spread backwards towards the stimulation site. We compared the nearest edge distances between the initial time point of depression onset and the end of depression (up to 10 s after the depression onset) among the different stimulation conditions (Fig. 7b). Even though the initial nearest edge distances varied among the stimulation conditions, the last nearest edge distances decreased in all conditions. Two-way ANOVA revealed significant differences in the nearest edge distances of the neural Ca^2+^ depression among the stimulation conditions and between the initial and final time points (stimulation condition, *F* = 4.80, *p* < 0.05; time point, *F* = 4.91, *p* < 0.05; stimulation condition × time point, *F* = 0.29, *p* = 0.94; post-hoc Tukey’s HSD test, 10-Hz ICMS vs visual stimulation, difference = 479 µm, *p* < 0.05; 25-Hz ICMS vs visual stimulation, difference = 526 µm, *p* < 0.05; 50-Hz ICMS vs visual stimulation, difference = 610 µm, *p* < 0.05; 100-Hz ICMS vs visual stimulation, difference = 613 µm, *p* < 0.05; initial vs last, difference = –168 µm, *p* < 0.05). However, the speeds of the nearest edge distance change did not significantly differ across stimulation conditions (Fig. 7c and d, one-way ANOVA, *F* = 0.70, *p* = 0.65).

**Figure 7.**
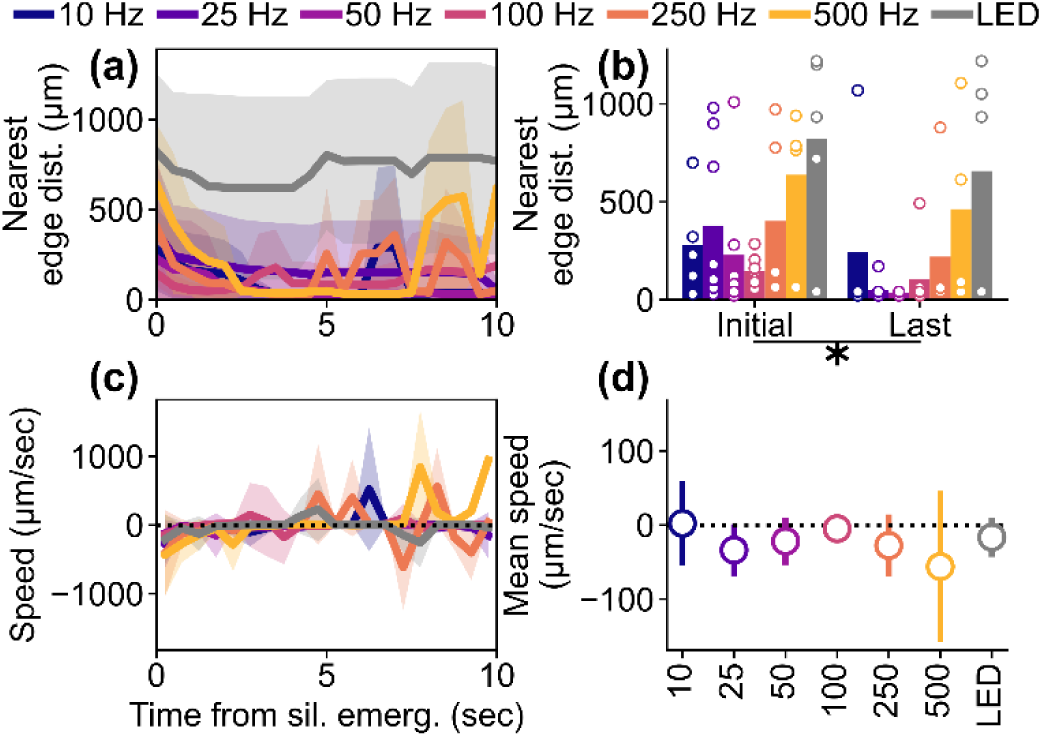
Dynamics of nearest ‘edge distance’ of depression. (a) Temporal change of the nearest ‘edge distance’ of the depression induced by the various stimulation conditions (mean ± 1SD among the animals). The data were aligned to the initiation of each depression. (b) Comparison of the initial and final nearest edge distances among the different stimulation conditions. (c) Dynamics of the speed of the nearest edge distance of depression, indicating movement away from the stimulation site (positive) or towards it (negative) (mean ± 1SD among the animals). (d) Averaged speed of nearest edge distance (mean ± 1SD among the animals). * indicates *p* < 0.05 for pairwise comparison with post-hoc Tukey’s HSD test.

The shrinking activation area at high frequencies may reflect a strong drive of inhibitory neurons, leading to transient and localized activation. In contrast, at low frequencies, the inhibitory drive remains low, and activation area propagates, possibly through orthodromic, post-synaptic activation of the network. This explanation is supported by the observed depression phenomena. The higher-frequency ICMS likely elicits stronger inhibition during the early time point, leaving little room for depression to occur after the activation ends. On the other hand, the lower frequency may engage inhibition gradually, enabling the depression to spread over the activation (see Discussion for frequency preference of inhibitory neurons). These findings regarding the spatiotemporal dynamics of stimulation-induced activation and depression provide valuable insights for determining the most effective stimulation train duration at different frequencies.

### Visual response is attenuated around the probe

The previous analysis revealed a tendency for weakened visually-evoked activation in the vicinity of the probe implanted in the visual cortex (Figs. 2g and 4h). To further investigate this reduced activation, we conducted additional analysis (Fig. S3). By comparing the induced activation and depression near the probe (27.5 µm < distance ≤ 150 µm, where 27.5 µm represents half of the probe shank width) and distant locations (150 µm < distance ≤ 500 µm), we observed clear reductions in responses near the probe (Fig. S3a and b). Pairwise comparisons with the pooled dataset confirmed that the responses near the probe were significantly lower than those in the distal regions during the transient offset period (10–12 s) (Fig. S3c and d; one-sided t-test, 0–2 s, *t* = –1.92, *p* = 0.06; 2–10 s, *t* = –1.95, *p* = 0.06; 10–12 s, *t* = –2.66, *p* < 0.05; 12–20 s, *t* = –0.65, *p* = 0.28). This reduction in response may reflect the tissue reactions associated with the probe implantation injury. Further investigations, especially focused on chronic degeneration and the recovery process of the visual response, are needed to elucidate the underlying mechanism.

### ICMS-induced metabolic hyper-supply lasts after Ca^2+^ activation

We next turned our attention to the relationship between Ca^2+^ activity and the hemodynamic response. Metabolically sensitive metrics such as oxyhemoglobin and deoxyhemoglobin signals play a crucial role in estimating energy supply and consumption to local neural networks [45], which are not yet fully understood in terms of the ICMS-induced responses. Similar to what we observed in neural Ca^2+^ signals (Fig. 4), we found that oxyhemoglobin and deoxyhemoglobin signals exhibited distinct spatiotemporal dynamics in response to different ICMS frequencies in a representative mouse (Fig. 8 and Movies S4–S9).

**Figure 8.**
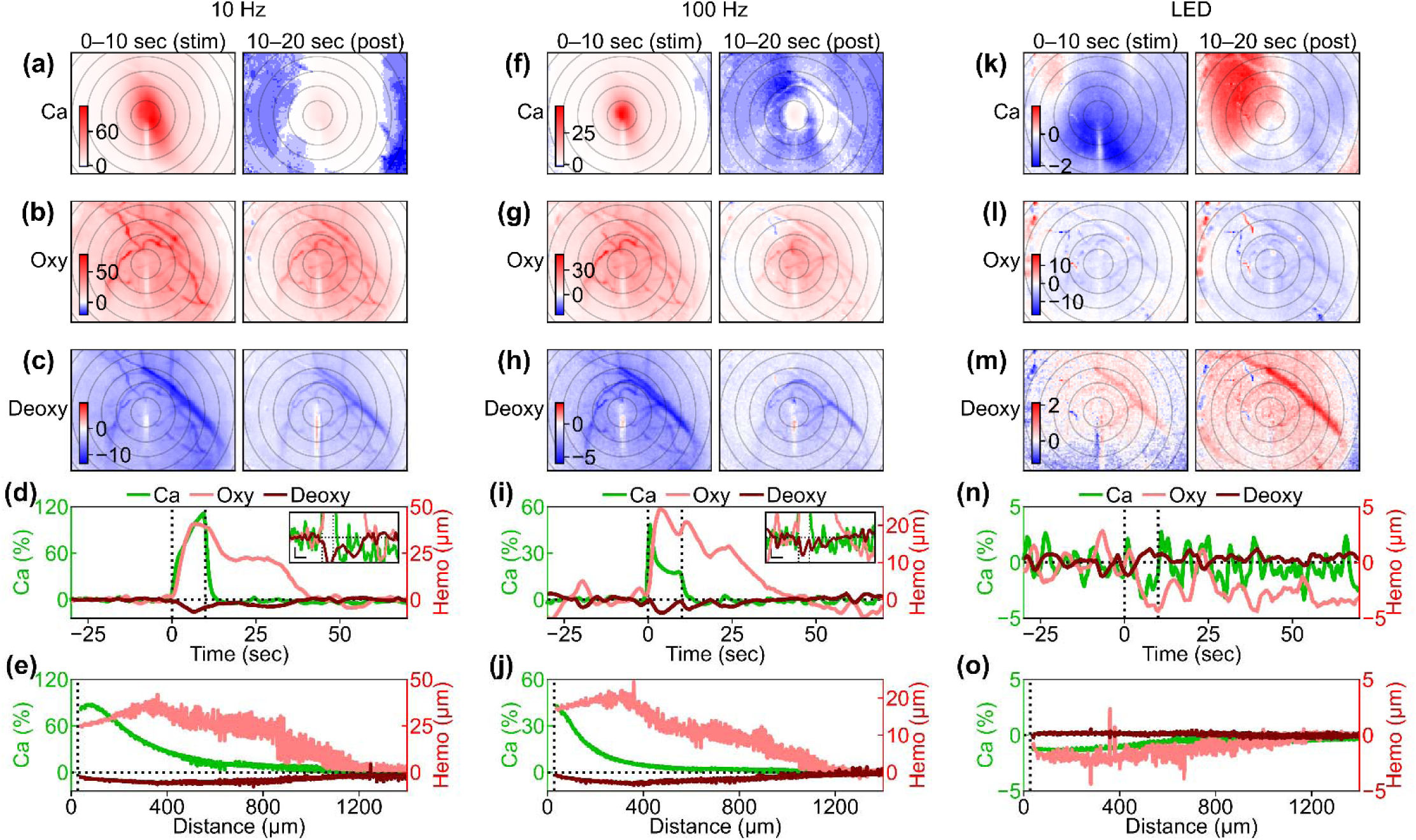
Spatiotemporal comparisons between the induced Ca^2+^ activation and hemodynamic responses. (a–c) Comparison of Ca^2+^, oxyhemoglobin, and deoxyhemoglobin responses to the 10-Hz ICMS during the stimulation period (0–10 s) and the post-stimulation period (10–20 s). Color scales are represented in % for Ca^2+^ response, µM for oxyhemoglobin, and µM for deoxyhemoglobin in (a), (b), and (c), respectively. Concentric circles indicate distances from the stimulation site, plotted at 200-µm intervals. (d) Time courses of the responses to the 10-Hz ICMS within 200 µm from the stimulation site. Inset indicates magnified time courses for better visibility (horizontal scale bar = 10 s; vertical scale bar = 2% and 2 µM for Ca^2+^ and hemodynamic signals, respectively). (e) Distance profiles of the responses to the 10-Hz ICMS. (f–j) Same metrics evaluated under 100-Hz ICMS condition. (k– o) Same metrics evaluated under visual stimulation condition.

During the 10-Hz ICMS train (0–10 s), the oxyhemoglobin concentration increased over a wider cortical area surrounding the stimulation site compared to the Ca^2+^ changes (Fig. 8b). This suggests that oxygen was metabolically supplied not only at the center of activation but also in the peripheral regions where clear excitatory Ca^2+^ activity was not observed. Even after the stimulation train ended (10–20 s), the oxyhemoglobin concentration remained elevated compared to the pre-stimulation baseline (increase in metabolic supply). With the 100-Hz ICMS (Fig. 8g), which induced more localized neural Ca^2+^ activation compared to the 10-Hz ICMS, the increase in oxyhemoglobin was confined to a narrower distance. Simultaneously, the deoxyhemoglobin showed a similar trend, decreasing in concentration (Fig. 8c and h; decrease in metabolic demand, relative increase in metabolic supply). It is worth noting that the changes in deoxyhemoglobin (< –10 µM in the 10-Hz ICMS and < –5 µM in the 100-Hz ICMS) were about five times smaller than those in oxyhemoglobin (> 50 µM and > 25 µM), indicating that the change in total hemoglobin concentration (sum of oxyhemoglobin and deoxyhemoglobin concentration changes, Δ[HbO] + Δ[HbR]) decreased over a wide cortical area during and after the stimulation trains. The hemodynamic signals emerged slightly slower but persisted substantially longer than the excitatory Ca^2+^ signals (Fig. 8d). Additionally, The spatial profile also revealed the evident difference between the Ca^2+^ and hemodynamic signals (Fig. 8e). Notably, the hemodynamic responses exhibited substantial changes over a wider area than the Ca^2+^ changes. This difference was attenuated with higher ICMS frequency (10-Hz ICMS v.s. 100-Hz ICMS). Compared to the ICMS-induced responses, visually-evoked hemodynamic responses were weaker (Fig. 8k–o).

We then quantified the distance, latency, and duration of the Ca^2+^ activation, oxyhemoglobin, and deoxyhemoglobin signals across all stimulation conditions (Fig. 9). The distance was significantly dependent on the stimulation conditions, but was independent of the signal modalities (Fig. 9a; two-way ANOVA, stimulation condition, *F* = 3.31, *p* < 0.05; signal modality, *F* = 0.92, *p* = 0.40; stimulation condition × signal modality, *F* = 0.52, *p* = 0.90). Post-hoc tests revealed the significant differences between several stimulation condition pairs (50-Hz ICMS vs 500-Hz ICMS, difference = –298 µm, *p* < 0.05; 500-Hz ICMS vs visual stimulation, difference = 326 µm, *p* < 0.05). Comparing the distances of the oxyhemoglobin and deoxyhemoglobin signals to the Ca^2+^ activation distance, we found that the oxyhemoglobin signal distances were 29 µm further than the Ca^2+^ signal, while the deoxyhemoglobin signal threshold distances were 73 µm further.

**Figure 9.**
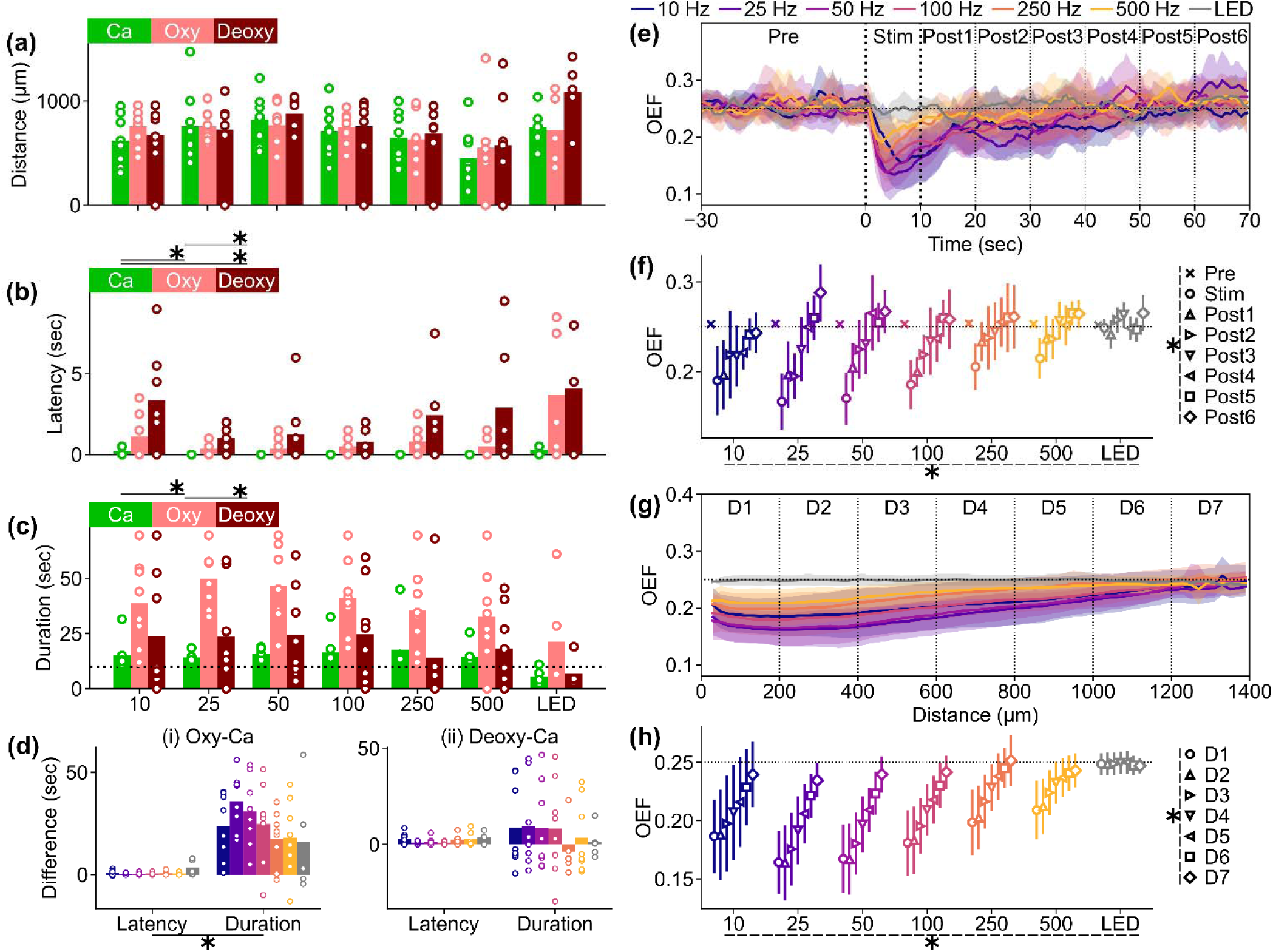
Comparisons of distances, latencies, and durations between excitatory neural Ca^2+^ and hemodynamic signals. (a) Distances of the induced Ca^2+^, oxyhemoglobin, and deoxyhemoglobin responses and comparisons across different stimulation conditions and signal modalities (Ca^2+^, oxyhemoglobin, and deoxyhemoglobin). (b) Latencies of the Ca^2+^ and hemodynamic signals. (c) Durations of the Ca^2+^ and hemodynamic signals. (d) Differences in latencies and durations between Ca^2+^ and oxyhemoglobin signals (i) and between Ca^2+^ and deoxyhemoglobin signals (ii) across different stimulation conditions. Higher positive values indicate tha the hemodynamic signals have longer latencies and durations compared to the Ca^2+^ signals. (e) Time courses of OEF in various stimulation conditions, which are divided into multiple groups based on timing (vertical lines). Lines and shaded areas indicate mean ± 1SD across mice. (f) Comparisons of estimated OEF across stimulation conditions and timing bins. Results of pairwise compa ison with post-hoc Tukey’s HSD test are summarized in Tables S6 and S7 for better visibility. (g) Distance profiles of OEF in various stimulation conditions (0–10 s), which are divided into multiple groups based on distance from the stimulation site (vertical dotted lines). Lines and shaded areas indicate mean ± 1SD across mice. (h) Comparisons of estimated OEF across stimulation conditions and distance bins. Results of pairwise comparison with post-hoc Tukey’s HSD test are summarized in Tables S8 and S9 for better visibility. * with solid line indicates *p* < 0.05 for post-hoc Tukey’s HSD test. * with dashed line indicates *p* < 0.05 for two-way ANOVA.

Significant differences in latencies were observed across the stimulation conditions and signal modalities (Fig. 9b; two-way ANOVA, stimulation condition, *F* = 3.95, *p* < 0.05; signal modality, *F* = 18.97, *p* < 0.05; stimulation condition × signal modality, *F* = 1.21, *p* = 0.28). Interestingly, the post-hoc Tukey’s HSD test revealed significant differences in latencies between certain ICMS frequencies and visual stimulation conditions (25-Hz ICMS vs visual stimulation, difference = 2 s, *p* < 0.05; 50-Hz ICMS vs visual stimulation, difference = 2 s, *p* < 0.05; 100-Hz ICMS vs visual stimulation, difference = 2 s, *p* < 0.05). This indicates that visual stimulation resulted in significantly slower response emergence compared to ICMS conditions, potentially due to the transmission of visual signals from the eye to the visual cortex and the unsynchronized recruitment of neurons, causing a sluggish emergence of signals. Furthermore, the latencies in the hemodynamic signals were a few seconds slower than the Ca^2+^ signals (oxyhemoglobin-Ca^2+^ = 1 s, *p* < 0.05; deoxyhemoglobin-Ca^2+^ = 2 s, *p* < 0.05). Additionally, the deoxyhemoglobin signal showed a significantly slower latency compared to the oxyhemoglobin signal (deoxyhemoglobin-oxyhemoglobin = 1 s, *p* < 0.05), which may reflect a cascade of blood influx and deoxygenation.

Having revealed distinct activation distance and latency differences in Ca^2+^ and hemodynamic activity dependent on stimulation parameters, we investigated whether there were differences in the duration of these signals during and following ICMS. Similar to the latencies mentioned earlier, the durations were significantly influenced by both the stimulation conditions and signal modalities (Fig. 9c; two-way ANOVA, stimulation condition, *F* = 2.51, *p* < 0.05; signal modality, *F* = 30.92, *p* < 0.05; stimulation condition × signal modality, *F* = 0.46, *p* = 0.94). The duration, particularly in oxyhemoglobin signal, tended to be longer around the 25-Hz ICMS condition (no significant differences in the post-hoc Tukey’s HSD test among the stimulation conditions). The gross durations across all stimulation conditions in the oxyhemoglobin signal were significantly longer compared to the Ca^2+^ and deoxyhemoglobin signals (post-hoc Tukey’s HSD test, oxyhemoglobin-Ca^2+^ = 24 s, *p* < 0.05; oxyhemoglobin-deoxyhemoglobin = 19 s, *p* < 0.05; deoxyhemoglobin-Ca^2+^ = 5 s, *p* = 0.23). This prolonged oxyhemoglobin signal was statistically significant (Fig. 9d, differences in latencies and durations between Ca^2+^ and hemodynamic signals; Ca^2+^ vs. oxyhemoglobin, *F*_stim_ = 1.15, *p*_stim_ = 0.34, *F*_time_ = 91.65, *p*_time_ < 0.05, *F*_stim × time_ = 1.46, *p*_stim × time_ = 0.20, post-hoc Tukey’s HSD test, latency vs duration, difference = 23 s, *p* < 0.05; Ca^2+^ vs. deoxyhemoglobin, *F*_stim_ = 0.40, *p*_stim_ = 0.88, *F*_time_ = 1.28, *p*_time_ = 0.26, *F*_stim × time_ = 0.54, *p*_stim × time_ = 0.77).

Therefore, although the degree of neural Ca^2+^ activity and the metabolic supply and demand in response to various ICMS frequencies and visual stimulation are highly dependent on the stimulation conditions, the prolonged increase in oxyhemoglobin concentration is consistently observed in most stimulation conditions. These results suggest that hemodynamic activity is involved in forming the post-activation Ca^2+^ response in excitatory neurons, in which the increased oxygenation may be associated with increased activity in inhibitory neurons and/or govern the depression phenomenon (see Discussion).

Alternatively, these prolonged hemodynamic signals could potentially reflect a metabolic supply-demand imbalance, leading to metabolic shortage and a reduction in neural activity (“metabolic silencing”), and/or indirectly depressing excitatory neurons through inhibitory neurons (see Discussion). To quantify the degree of metabolic demand during and after the stimulation within the range showing clear hemodynamic responses (< 500 µm, Figure 9e and f), we calculated the OEF (oxygen extraction fraction; Equation 5) with several baseline concentration assumptions (see Methods). OEF is defined as the fraction of deoxyhemoglobin concentration over the total hemoglobin concentration. We observed a significant reduction in OEF, particularly with the 10-, 25-, and 100-Hz ICMS during the stimulation and initial 20-s post-stimulation periods (two-way ANOVA, stimulation condition, *F* = 8.27, *p* < 0.05, timing, F = 36.45, *p* < 0.05, stimulation condition × timing, *F* = 1.69, *p* < 0.05; post-hoc Tukey’s HSD test, 10-Hz ICMS vs 250-Hz ICMS, difference = 0.021, *p* < 0.05, 10-Hz ICMS vs 500-Hz ICMS, difference = 0.025, *p* < 0.05, 10-Hz ICMS vs LED, difference = 0.030, *p* < 0.05, 25-Hz ICMS vs LED, difference = 0.024, *p* < 0.05, 100-Hz ICMS vs LED, difference = 0.022, *p* < 0.05; see Table S2 for results of post-hoc Tukey’s HSD test across the timing). These findings suggest that the 25–50-Hz ICMS trigger hyperoxygenation in the local tissues.

Additionally, we also asked how the induced hyperoxygenation is localized in the visual cortex (Fig. 9g). The OEF distance profiles during the 0–10-s stimulation period showed extensive spreads, nearly encompassing the entire imaging field upon ICMS application. Additionally, higher-frequency ICMS resulted in a diminished reduction of OEF. Regarding visual stimulation, the altered OEF is not discernible at any distance. Statistical quantification (Fig. 9h) revealed significant OEF reductions with 10–250 Hz ICMS (two-way ANOVA, stimulation condition, *F* = 24.12, *p* < 0.05; see Table S3 for results of post-hoc Tukey’s HSD test across stimulation conditions). These large reductions of OEF were restricted within 1000 µm from the stimulation sites (two-way ANOVA, distance, *F* = 30.16, *p* < 0.05; see Table S4 for results of post-hoc Tukey’s HSD test across distances). No significant interaction between the stimulation conditions and distances was observed (*F* = 0.89, *p* = 0.65). Taken together, these spatiotemporal profiles suggest that the ICMS-induced change in OEF meets the metabolic demand that is caused by the natural sensory stimulus (visual stimulation), which is enhanced especially by low or middle frequencies.

Further analyses revealed that the correlation between spatiotemporal dynamics of excitatory neural Ca^2+^ responses and oxyhemoglobin responses during the stimulation period (0–10 s) significantly depended on the stimulation conditions (Fig. S4a, one-way ANOVA, *F* = 3.02, *p* < 0.05). Specifically, the Ca^2+^ responses in the lower ICMS frequencies, particularly at 10 Hz, tended to be positively correlated with oxyhemoglobin responses during the stimulation train period (post-hoc Tukey’s HSD test, 10-Hz ICMS vs 500-Hz ICMS, difference = –0.31, *p* < 0.05). Furthermore, strong correlations were observed in the magnitudes extracted from the Ca^2+^ activation/depression and oxy/deoxyhemoglobin signals (Fig. S4e–h, Spearman’s rank correlation coefficient, *p* < 0.05). This suggests that the 10– and 100-Hz ICMS induced a positive coupling between neural activity and oxygen supply/demand, whereas visual stimulation resulted in a negative coupling between neural depression and oxygen demand.

### Soma and neuropil exhibit frequency-dependent ICMS-induced activation and depression

We investigated the relationship between stimulation parameters and the activation and depression of excitatory neurons, as well as oxyhemoglobin and deoxyhemoglobin signals, using mesoscopic-scale widefield imaging. Although this imaging technique allowed us to capture spatial information across a large area of the visual cortex, it cannot differentiate signals in finer neural structures like soma and neuropil. The mechanisms of depression in the soma and neuropil are not well understood. To address this, we conducted two-photon imaging (Fig. 10 and Movies S10–S12).

**Figure 10.**
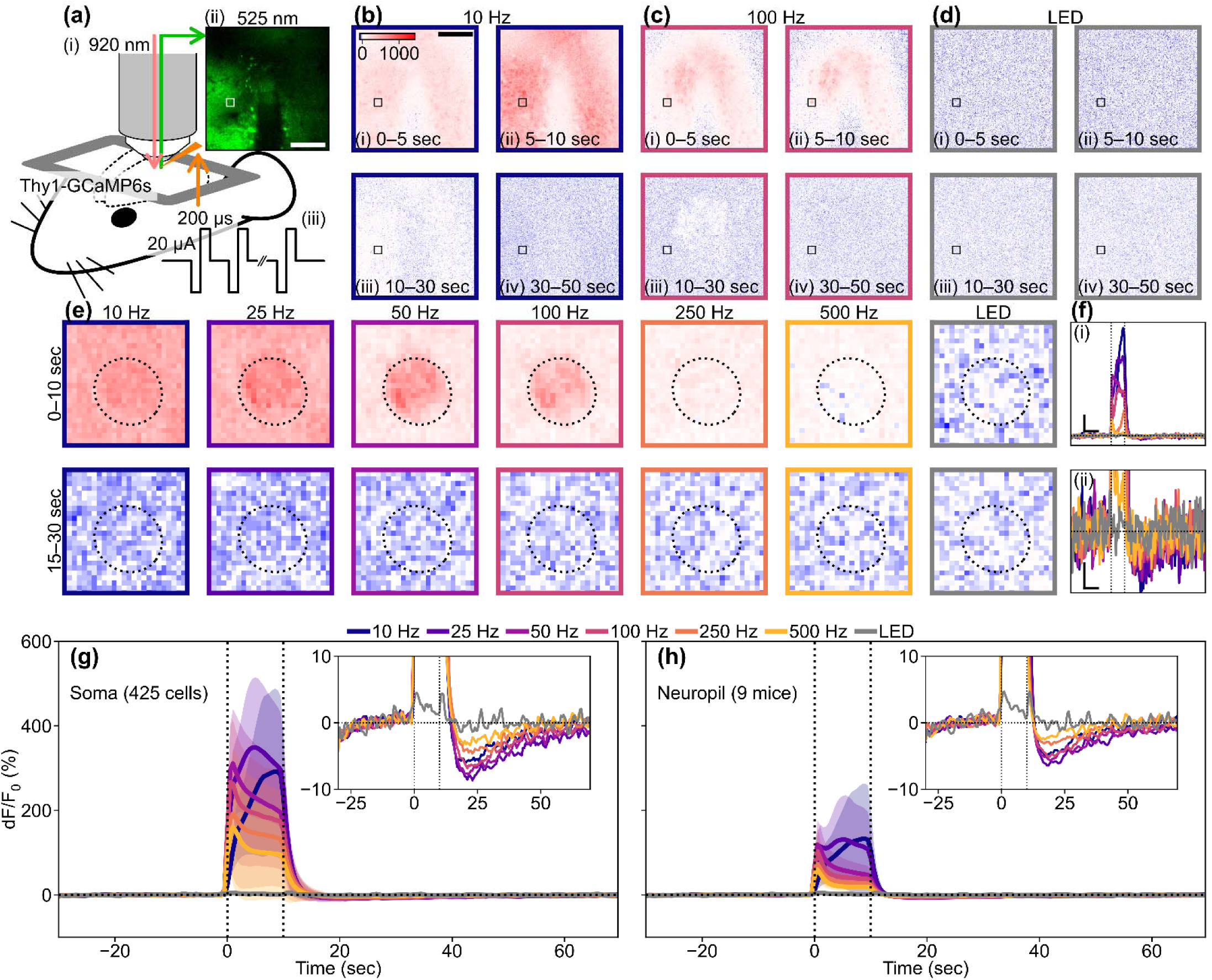
Two-photon data quantifying somatic and neuropil responses induced by ICMS and visual stimulation. (a) Experimental setup. Infrared two-photon laser was applied to Thy1-GCaMP6s mouse (i) and a green fluorescent image was acquired (ii) before, during, and after the stimulation train period (iii). Scale bar = 100 µm. (b–d) Time-lapse images showing the responses to (b) 10-ICMS, (c) 100-Hz ICMS, and (d) visual stimulation. Scale bar = 100 µm. The color scale unit is %, and the same color scales are shared in (e). (e) Somatic Ca^2+^ responses of an example neuron (white and black boxes in a–d) to the various stimulation conditions. Dotted ellipses indicate the ROI of the neuron. (f) Time courses of the somatic Ca^2+^ responses shown in (d). Line colors correspond to panel colors in (b)–(e). Vertical scale bars in (i) and (ii) = 100% and 10%, respectively. Horizontal scale bars in (i) and (ii) = 10 sec. Vertical dotted lines indicate the onset and offset of the stimulation train. (g and h) Mean ± 1SD of somatic (g, n = 425 cells) and neuropil (h, n = 9 mice) Ca^2+^ time courses. Colors correspond to the stimulation conditions. Vertical dotted lines indicate the onset and offset of the stimulation train. The inset shows mean time courses with a magnified y scale to clearly show the depression.

Under two-photon microscopy, we observed strong scattered spots of somatic autofluorescence around the inserted probe and neurite blebs (Fig. 10aii). Despite these autofluorescent spots, we were able to observe both somatic and neuropil activities during and after the 10-Hz ICMS train (Fig. 10b). The somas exhibited relatively stronger activation compared to the neuropil. In the 100-Hz ICMS condition, the somas near the stimulation site showed sparse activation, and the neuropil activation was localized near the stimulation site (Fig. 10c). In both cases, we observed distinct depression in both the somas and neuropil, where the Ca^2+^ activity dipped below the baseline resting state before the stimulus. We manually defined the ROI for each neuron (see Methods) and obtained somatic Ca^2+^ time courses by averaging dF/F_0_ values inside single ROIs. We also obtained neuropil Ca^2+^ time courses by averaging dF/F_0_ values outside the ROIs.

To examine the dependency of excitatory neural Ca^2+^ response on the stimulation condition, we selected one ROI as an example (Fig. 10e and f). These somatic and surrounding neuropil responses were dependent on the stimulation conditions (Fig. 10e).

For instance, during the 10-s stimulation trains, the ICMS-induced Ca^2+^ activity was highest at 10 Hz both in the soma and neuropil, and decreased as the ICMS frequency increased (Fig. 10e, top, 0–10 s). This observation aligns with the results we obtained using mesoscopic-scale microscopy (Figs. 2 and 4). A few seconds after the 10-s stimulation train ended, dominant depression was observed in both the soma and neuropil (Fig. 10e, bottom, 15–30 s). With cellular scale imaging, we were able to discern the visual stimulation preference of each neuron for the stimulation onset, offset, and onset/offset (Fig. 10d, Movie S12). Although the visual responses were remarkably weaker than the ICMS responses, similar to the mesoscopic-scale imaging findings, several neurons exhibited Ca^2+^ activation at different time points during the stimulation train.

The somatic Ca^2+^ signals (Fig. 10fi) exhibited a partially similar time course compared to the results obtained from mesoscopic-scale imaging (Fig. 4i). In mesoscopic-scale imaging, lower-frequency ICMS resulted in ‘developing activation’ (activation that slowly increases with stimulus duration), while higher-frequency ICMS caused a transient one. In this example soma (Fig 10), the higher-frequency ICMS initially induced a transient activation, followed by developing activation. With the exception for the visual stimulation condition, we consistently observed distinct somatic depression following activation (Fig. 10fii). We analyzed the neural Ca^2+^ time courses of 425 somatic ROIs and of neuropil in 9 mice (Fig. 10g and h). Overall, the pooled dataset revealed developing activation with lower-frequency ICMS, whereas higher-frequency ICMS triggered an initial transient increase in Ca^2+^ fluorescent intensity, followed by rapid reduction. However, all ICMS conditions maintained activation throughout the 10-s stimulation train. The depression phenomena were also observable in all ICMS conditions, regardless of the different magnitudes. Additionally, we observed weak transient on/off activation with a visual stimulation train. These trends were consistent in neuropils, although the activation magnitudes were noticeably smaller compared to somas. We compared the somatic activation magnitudes between the transient (0–2 s) and sustained (2–10 s) phases of the stimulation trains (Fig. S5a and b), and found that the transient-dominant responses were evident in the 50–500 Hz ICMS and visual stimulation conditions (50 Hz ICMS, paired t-test, *t* = 12.28, *p* < 0.05; 100 Hz, *t* = 13.62, *p* < 0.05; 250 Hz, *t* = 6.95, *p* < 0.05; 500 Hz, *t* = 10.88, *p* < 0.05; visual stimulation, *t* = 7.98, *p* < 0.05). On the other hand, the sustained-dominant responses were observed in the 10– and 25-Hz ICMS conditions (10 Hz, *t* = –28.92, *p* < 0.05; 25 Hz, *t* = –21.98, *p* < 0.05). Moreover, the change rates of somatic Ca^2+^ response magnitude during the stimulation train significantly varied among the stimulation conditions (Fig. S5c and d, one-way ANOVA, *F* = 395.77, *p* < 0.05). Lower ICMS frequencies tended to induce an increasing activation, while higher-frequency ICMS and visual stimulation tended to induce relatively constant activation (Table S1). Overall, these findings demonstrate that the somas and neuropils identified through two-photon microscopy exhibited response properties similar to those observed under mesoscopic-scale microscopy.

Given that the spatial extent of the mesoscopic-scale Ca^2+^ responses differed across ICMS frequencies (Fig. 4), we investigated how the response magnitude of somas at different distances from the stimulation site was influenced by ICMS frequencies (Fig. S6). Firstly, we quantified the distribution of distances for the activated somatic ROIs (Fig. S6a). The identified somas were mostly located around 50–100 µm from the center of the stimulation channel. Since the probe had a width of 55 µm, tissues within this range could be deformed, and might have displaced neurons. Consistent with the mesoscopic-scale results, the magnitudes of somatic activation and depression were dependent on distance and stimulation conditions (Fig. S6b, activation; two-way ANOVA, distance, *F* = 52.29, *p* < 0.05; stimulation condition, *F* = 334.34, *p* < 0.05; Figure S6c, depression; distance, *F* = 5.14, *p* < 0.05; stimulation condition, *F* = 17.62, *p* < 0.05). Further analyses revealed significant differences in magnitudes between several distances for activation (post-hoc Tukey’s HSD test, 0– 50 µm vs 50–100 µm, difference = 52.24%, *p* < 0.05; 0–50 µm vs 150–200 µm, difference = –27.51%, *p* < 0.05; 0–50 µm vs 200–250 µm, difference = –41.91 µm, *p* < 0.05; 50–100 µm vs 100–150 µm, difference = –49.04%, *p* < 0.05; 50–100 µm vs 150–200 µm, difference = –79.75%, *p* < 0.05; 50–100 µm vs 200–250 µm, difference = –94.15%, *p* < 0.05; 50–100 µm vs 250–300 µm, difference = –112.77%, *p* < 0.05; 100–150 µm vs 150–200 µm, difference = –30.72%, *p* < 0.05; 100–150 µm vs 200–250 µm, difference = –45.12%, *p* < 0.05) and for depression (100–150 µm vs 150–200 µm, difference = –0.57%, *p* < 0.05; 100–150 µm vs 200–250 µm, difference = –0.63%, *p* < 0.05). Moreover, the interaction between stimulation condition and distance significantly affected the magnitudes of somatic activation and depression (two-way ANOVA, activation, *F* = 6.01, *p* < 0.05; depression, *F* = 2.24, *p* < 0.05), suggesting that ICMSs at different frequencies could activate/depress somas at different distances. These results demonstrate that somatic Ca^2+^ responses, akin to mesoscopic-scale responses (Fig. S7), preferred specific ICMS frequencies corresponding to the distance from the stimulation site.

As demonstrated, Ca^2+^ responses in soma and neuropil of excitatory neurons were influenced by stimulation conditions. Thus, we proceeded to statistically compare the magnitude and duration of activation and depression in both somas and neuropils, as well as their balance (ratios of magnitudes between soma and neuropil, and between activation and depression) across different stimulation conditions (Fig. 11a).

**Figure 11.**
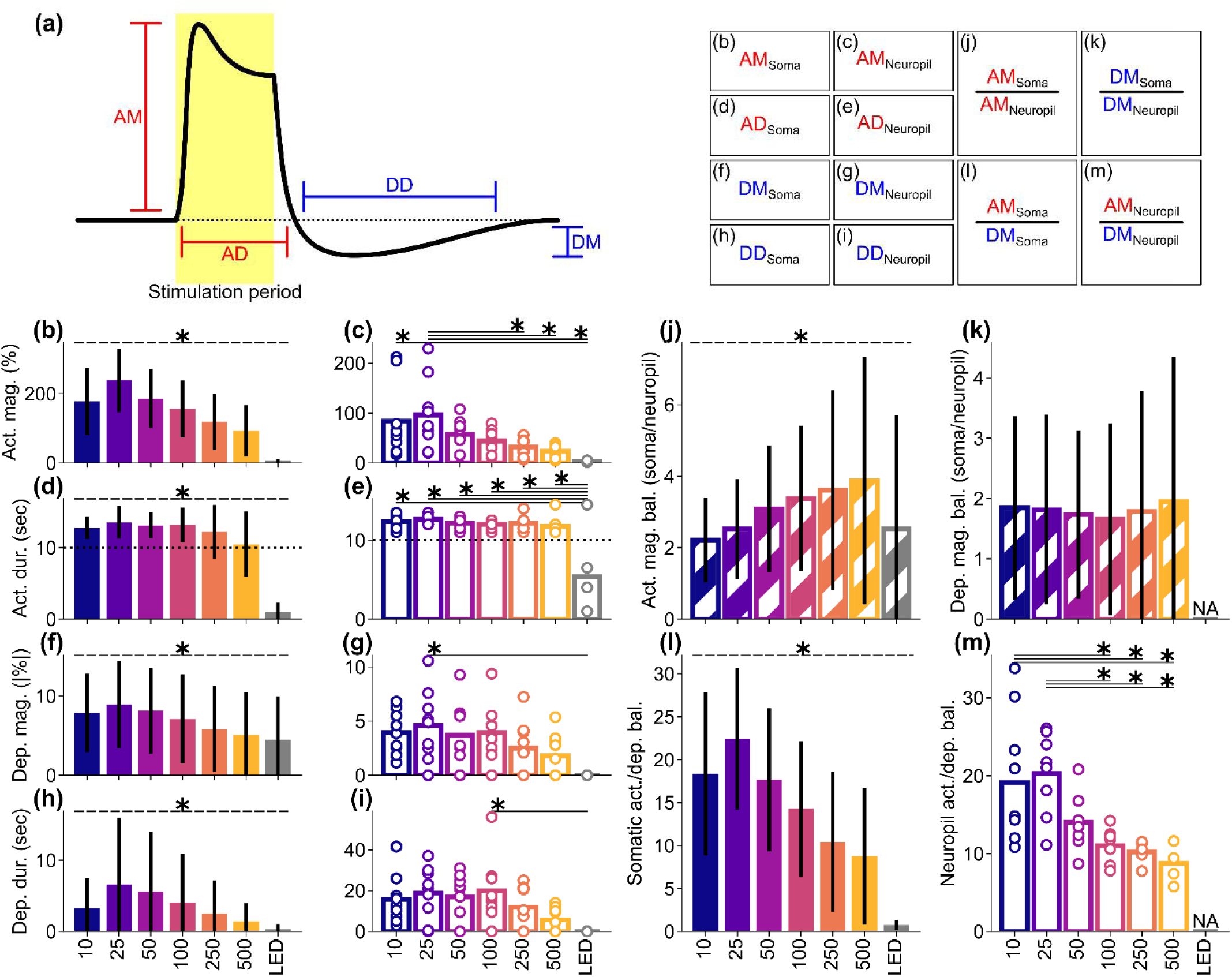
Summary of somatic and neuropil activation and depression observed under two-photon microscopy. (a) (left) Schematic representation of the measured activation and depression metrics. Activation magnitude (AM; b and c), activation duration (AD; d and e), depression magnitude (DM; f and g), and depression duration (DD; h and i) were extracted from the Ca^2+^ response time course in somatic ROIs and neuropil. (right) Description of the measurements and calculation, corresponding to the panels below (b–m). (b, d, f, and h) Somatic metrics: (b) activation magnitude, (d) activation duration, (f) depression magnitude, (h) depression duration. (c, e, g, and i) Neuropil metrics. (j) Activation magnitude balance between soma and neuropil. A higher value (> 1) indicates stronger activation in the soma compared to the neuropil. (k) Depression magnitude balance between soma and neuropil. A higher value (> 1) indicates stronger depression in the soma compared to the neuropil. (l) Somatic activation/depression balance. A higher value (> 1) indicates stronger activation compared to depression. (m) Neuropil activation/depression balance. A higher value (> 1) indicates stronger activation compared to depression. * with horizontal dashed line indicates *p* < 0.05 for one-way ANOVA on the somatic metrics (b, d, e, h, and j–l). Results of pairwise comparison with post-hoc Tukey’s HSD test are summarized in Tables S7–12 for better visibility of these panels. * with horizontal solid line indicates *p* < 0.05 for pairwise comparison with post-hoc Tukey’s HSD test (c, e, g, i, and m). If samples showed no significant activation and/or depression, their metrics were replaced with zero instead of being excluded from analyses for completing statistical evaluation (b–i). If the denominators of the balance metrics were zero, these balance metrics were excluded (j–m).

The magnitudes of somatic activation were significantly affected by the stimulation conditions (Fig. 11b, one-way ANOVA, *F* = 280.43, *p* < 0.05), peaking at the 25-Hz ICMS (see Table S5 for pairwise comparisons with post-hoc Tukey’s HSD test). In contrast, the duration of activation exhibited a relatively uniform distribution, except under the visual stimulation condition (Fig. 11d, *F* = 697.07, *p* < 0.05; Table S6). Similarly, the magnitudes of the somatic depression were significantly influenced by the stimulation conditions, displaying an unimodal distribution biased towards 25 Hz (Fig. 11f, *F* = 35.17, *p* < 0.05; Table S7). However, the distribution of somatic depression magnitudes was broader than that of the activation.

Consequently, the balance between somatic activation and depression (activation magnitude/depression magnitude) exhibited an unimodal preference, with a peak at the 25-Hz ICMS condition (Fig. 11l, *F* = 147.35, *p* < 0.05; Table S10). This indicates that lower-frequency ICMS, particularly at 25 Hz, elicited activation-dominant somatic Ca^2+^ responses. Interestingly, the activation/depression balance in the visual stimulation condition was nearly one, which suggests that visual stimulation induced a balanced activation of excitatory and inhibitory/suppressive responses. To replicate neural responses induced by visual stimulation, higher-frequency ICMS could be utilized in the field of visual cortical prostheses (see Discussion). Moreover, the duration of somatic depression also exhibited a clear bias toward the 25-Hz ICMS condition (Fig. 11h, *F* = 47.01, *p* < 0.05; Table S8), and tended to be shorter than the duration of activation.

Given the clear dependency of somatic activation and depression on ICMS frequencies and visual stimulation, we proceeded to quantify the responsive properties of neuropil across various stimulation conditions and compared them with somatic properties. The magnitudes of activation in the neuropil displayed a preference similar to that observed in somas (Fig. 11c), was significantly influenced by the stimulation conditions (one-way ANOVA, *F* = 4.42, *p* < 0.05), and exhibited a unimodal distribution peaking at the 25-Hz ICMS (post-hoc Tukey’s HSD test, 10-Hz ICMS vs visual stimulation, difference = –79.78%, *p* < 0.05; 25-Hz ICMS vs 250-Hz ICMS, difference = –64.53%, *p* < 0.05; 25-Hz ICMS vs 500-Hz ICMS, difference = –72.61%, *p* < 0.05; 25-Hz ICMS vs visual stimulation, difference = –92.38%, *p* < 0.05).

When comparing the magnitudes of activation between soma and neuropil, we observed a significant difference in the soma/neuropil balance across stimulation conditions (Fig. 11j, *F* = 28.99, *p* < 0.05; Table S9), where higher-frequency ICMS elicited more soma-dominant activations. The duration of activation in the neuropil exhibited a uniform distribution except for the visual stimulation, similar to somas (Fig. 11e, *F* = 11.88, *p* < 0.05), in which visual stimulation induced significantly shorter activation compared to all ICMS frequencies (10-Hz ICMS vs visual stimulation, difference = –6.93 s, *p* < 0.05; 25-Hz ICMS vs visual stimulation, difference = –7.21 s, *p* < 0.05; 50-Hz ICMS vs visual stimulation, difference = –6.77 s, *p* < 0.05; 100-Hz ICMS vs visual stimulation, difference = –6.60 s, *p* < 0.05; 250-Hz ICMS vs visual stimulation, difference = –6.71 s, *p* < 0.05; 500-Hz ICMS vs visual stimulation, difference = –6.38 s, *p* < 0.05). These results indicate that both somas and neuropils are capable of sustained activation throughout the ICMS train, at least for a duration of 10 s, within 10–500 Hz, although the magnitudes may vary depending on the frequency.

The depression magnitude in the neuropil also exhibited a slight bias, peaking at 25 Hz (Fig. 11g, one-way ANOVA, *F* = 2.72, *p* < 0.05; post-hoc Tukey’s HSD test, 25-Hz ICMS vs visual stimulation, difference = –4.58%, *p* < 0.05). However, the visual stimulation did not induce significant depression in the neuropil. When comparing the depression magnitudes between somas and neuropils, we observed that the soma/neuropil balance of depression magnitudes were unaffected by the ICMS frequencies (Fig. 11k, *F* = 1.18, *p* = 0.32).

This suggests that the mechanism underlying the depression phenomenon selectively affects the soma and its selectivity is not modified by the frequent repetition of the ICMS pulses. The activation/depression balance in the neuropil also peaked at the 25-Hz ICMS condition (Fig. 11m, *F* = 6.71, *p* < 0.05; 10-Hz ICMS vs 100-Hz ICMS, difference = –8.10, *p* < 0.05; 10-Hz ICMS vs 250-Hz ICMS, difference = –8.90, *p* < 0.05; 10-Hz ICMS vs 500-Hz ICMS, difference = –10.40, *p* < 0.05; 25-Hz ICMS vs 100-Hz ICMS, difference = –9.27, *p* < 0.05; 25-Hz ICMS vs 250-Hz ICMS, difference = –10.07, *p* < 0.05; 25-Hz ICMS vs 500-Hz ICMS, difference = –11.57, *p* < 0.05). This finding is similar to somatic activation, suggesting that the 25-Hz ICMS could be the preferred choice for inducing an activation-dominant neural response in the field of neuroprosthetics. The depression duration in the neuropil exhibited a peak at around 100 Hz (Fig. 11i, *F* = 2.98, *p* < 0.05; post-hoc Tukey’s HSD test, 100-Hz ICMS vs visual stimulation, difference = –19.94 s, *p* < 0.05), which tended to be longer than in the somas. These results demonstrate that ICMS alone has the capacity to elicit depression, even in the neuropil, implying that ICMS and visual stimulation do not share the same mechanism for inducing the depression phenomenon.

Overall, the activation magnitudes exhibited a bias towards 25 Hz, whereas the depression magnitudes tended to be higher in higher frequencies. This discrepancy indicates that the mechanism underlying depression differs from that of activation, which shows a preference for higher-frequency ICMS.

## Discussion

In the current study, we showed that ICMS induced both activation and subsequent depression in neuronal Ca^2+^ signals from excitatory neurons in the visual cortex of head-restricted mice (Fig. 1). The magnitude and duration of activation and depression were found to be dependent on the frequency and modality of the stimulation (Figs. 2–7). Furthermore, the hemodynamic signals were significantly influenced by both the various-frequency ICMS and visual stimulation (Figs. 8 and 9). Through cellular-level two-photon imaging, we observed that both soma and neuropil exhibited dependency on the stimulation conditions, with depression occurring after the activation (Figs. 10 and 11). The distinct preferences of ICMS frequency and stimulation modality on induced activation and depression, as well as prolonged oxygenation, raise intriguing questions regarding the underlying mechanisms of the depression phenomenon. Potential factors to consider include inhibitory neurons and metabolic contributions among others. Exploring these mechanisms could provide valuable insights into the neurovascular coupling, inspire the development of biomimetic visual prostheses, and guide future directions in neurophysiological studies involving the ICMS.

### Possible mechanisms inducing the depression of the Ca^2+^ signals in the excitatory neurons

We observed that the depression of excitatory Ca^2+^ signals occurred following the activation (Fig. 2), and its characteristics varied depending on the stimulation parameters (Figs. 4 and 5). Similar depression phenomena, where neural activity is reduced compared to the pre-stimulation baseline, have been described in previous studies [6, 29, 31, 32], where the specific methods used for induction and quantification varied each other but the underlying mechanisms might be shared. For instance, McCreery and colleagues investigated a phenomenon called stimulation-induced depression of neuronal excitability (SIDNE) [32, 46, 47]. In their experiments, repetitive electrical pulses (100–500 Hz, < 3 nC/phase) were applied for 7 hours. They observed an increase in the threshold currents required to evoke compound action potentials after the pulse train. This SIDNE phenomenon could be reversed within several days, and no histological tissue injury attributable to the stimulation was observed. The mechanism underlying SIDNE is thought to be related to the influx of Ca^2+^, triggering intracellular mechanics affecting membrane ion channels [32], and/or transient changes of extracellular Ca^2+^ and potassium ion concentrations induced by prolonged stimulation [48].

Cortical spreading depression (CSD) is another example of an induced neural depression phenomenon, which can be induced by various forms of stimulation, including electrical pulses [49, 50]. CSD has often been investigated using hemodynamic signal imaging [51], where CSD is accompanied by a reduction in reflectance (indicating increased blood flow). Unlike SIDNE, CSD occurs on a shorter timescale, emerging a few seconds after the stimulation and lasting from several minutes [52–54] to an hour [55]. The formation of CSD involves both presynaptic and postsynaptic mechanisms. Presynaptically, there is a reduction in the probability of transmitter release, while postsynaptically there is a relative increase in inhibitory current, contributing to the generation of CSD [55].

In other scenarios, there have been reports of neural activity reduction occurring on an even shorter timescale, lasting in the range of hundreds of milliseconds. In these cases, the gamma-aminobutyric acid B (GABAb) receptor has been found to play a role in mediating the reduction of action potentials [31] and population-level membrane potential [6] following transient activation induced by ICMS. Furthermore, numerous other potential mechanisms can contribute to decreased neural excitability and related phenomena. These include energy supply shortage due to repetitive neural activation (metabolic limitation), neurotransmitter vesicle depletion, network inhibition involving vascular-astrocyte-neural coupling, network inhibition through recurrent inhibition (excitatory neuron – inhibitory neuron – excitatory neuron feedback loop), long-term depression, contrast enhancement through lateral inhibition, and sensory adaptation. When summarizing these possible mechanisms underlying the Ca^2+^ depression phenomenon, they can be classified into several groups based on different aspects such as spatial and temporal considerations.

#### Inhibitory circuits on spatiotemporal inhibition and depression

The depression exhibited a preference for higher-frequency ICMS compared to activation (Figs. 5 and 11). Unlike SIDNE, which is induced by long-duration ICMS train (e.g., 7 hr, [32]) leading to long-lasting neural excitability reduction (e.g., several days), our study utilized relatively short train duration (10 s) that caused depression of Ca^2+^ signals persistent for several tens of seconds. Previous literatures have also reported similar depression phenomena observed in various scales and methods, induced by various forms of exogenous stimulation, including ICMS [6, 29, 31, 56]. For example, Butovas et al. electrophysiologically observed a reduction in firing rates in the primary somatosensory cortex of anesthetized mice following ICMS pulse application [31]. Miyamoto et al. used voltage-sensitive dye imaging in the anesthetized rat visual cortex and found decreases in the fluorescent change ratio (dF/F_0_) after ICMS-induced increases [6]. These ICMS-induced depression phenomena were abolished by the application of the GABAb receptor antagonist (CGP46381) in both studies. In our study, we observed a slight shift in the frequency preference of the depression towards higher-frequency ICMS, indicating the capacity of inhibitory neurons to respond to higher frequency inputs and stimulation [57, 58]. These findings suggest that the depression of excitatory Ca^2+^ signals is mediated by inhibitory neuronal contributions, particularly through the involvement of GABAb receptors. This can involve both recurrent inhibition within local neural circuits and larger-scale networks encompassing non-neuronal components, like astrocytes [59]. Selective recording of inhibitory neural activity and/or controlling it in conjunction with excitatory Ca^2+^ imaging will provide valuable insights for further elucidating this aspect.

#### Metabolic stress and fatigue on neuronal depression and decrease in GCaMP signal

In the current study, we employed metabolic-sensitive imaging and observed a prolonged oxyhemoglobin increase, leading to a reduction in the relative oxygen consumption (OEF), during the Ca^2+^ depression period (Figs. 8 and 9). Our recording method for neural activity only captured Ca^2+^ signals in the excitatory neurons due to the use of Thy1-GCaMP6s mice (see Methods). We did not investigate inhibitory neural responses to the repetitive stimulation in this study. The prolonged changes in the oxyhemoglobin signal may reflect the extended or delayed inhibitory neural activity induced by the ICMS train, which plays a dominant role during the post-activation period of excitatory neurons. However, the selective activation of parvalbumin-positive interneurons reduces oxygenation [60]. Thus, the co-activation both of excitatory and inhibitory neurons may trigger the prolonged oxygenation observed in the current study.

There are additional factors that could contribute to the depression, such as metabolic exhaustion, depletion of cellular energy/oxygen supply, and synaptic fatigue [61]. It is important to note that intrinsic hemodynamic signal imaging can reflect broad changes in neuronal activity, not limited to specific temporal or local neural circuits. Even if there is an increase in local oxyhemoglobin concentration, it is unclear if the neurons in the same area consume the same amount of oxygen [62]. However, the ICMS induced a more substantial reduction in OEF compared to the visual stimulation (natural sensory input). This suggests that by applying ICMS, the local cortical tissues received oxygen-rich blood flow, mitigating the likelihood of metabolic exhaustion.

Synaptic fatigue can occur, especially with repetitive stimulation [63], as neurotransmitters released from excitatory neurons become depleted, leading to a suppressive effect. While this explanation suggests that the depression exhibits properties similar to activation in the surrounding excitatory neurons, our observations indicate that the depression displays a different ICMS frequency preference compared to activation (10 Hz vs 25 Hz in Figure 5e and f for activation and depression optimal frequencies under mesoscopic-scale imaging, respectively; activation/depression balance varying across ICMS frequencies in Figure 11l and m). Astrocytes also modulate (reduce) neurotransmitter uptake rate as the firing rate increases [64]. Therefore, synaptic fatigue may only partially contribute to the formation of depression.

There are likely multiple mechanisms underlying the excitatory neural Ca^2+^ dynamics induced by ICMS. One possible mechanism involves rapid inhibitory neural input from local neural circuits [65, 66], occurring in the initial period of the stimulation train (0–2 s) and/or in response to each stimulation pulse. This type of inhibition may also contribute to shaping the transient activation observed in higher-frequency ICMS conditions. A second mechanism involves network-level inhibition, which includes multiple-step excitatory-inhibitory connections and vasculo-glio-neural coupling [67–69]. This mechanism operates with longer latency and duration following activation compared to the local circuits. The repetitive stimulation used in our study may also induce another type of modulation, such as synaptic fatigue that reduces transmitter release probability [70], Ca^2+^ depletion in the extracellular space [71], potassium accumulation in the intracellular space [72], and synaptic plasticity that diminishes long-term transmission efficacy [73]. To further understand the contribution of these possible mechanisms to ICMS-induced response, it would be beneficial to control hemodynamic activity in a timing-dependent manner [74–76], visualize transmitter release [77, 78], and conduct longer post-stimulation imaging.

#### Inhibition in visual processing

The depression phenomena in the current study were observed using both ICMS and visual stimulation across multiple scales (mesoscopic and microscopic) and tissue categories (soma and neuropil). Notably, the depression induced by ICMS was weaker than the activation, while the activation and depression induced by visual stimulation showed balanced magnitudes (Figs. 5 and 11). Regarding activation, visual cortical tissues only exhibited transient activation in response to the 10-s train of visual stimulation. These differences strongly suggest that the underlying mechanisms responsible for depression in ICMS and visual stimulation are distinct. For the ICMS, the lower-frequency one, particularly at 25 Hz, elicited activation-dominant Ca^2+^ responses in soma. Therefore, if an activation, rather than depression, in somas is desired for neuroprosthetic application, the 25-Hz ICMS train would be the primary choice.

For the visual stimulation, we employed full-field illumination, which simultaneously activates all retinal neurons and recruits both excitatory neural projections and lateral inhibition [79]. Lateral inhibition normally enhances spatial contrast and reduces redundant information. The visual information is then relayed through the lateral geniculate nucleus and further processed in the visual cortex, where local neural networks apply additional lateral inhibitory processes [80, 81], that lead to reductions in neural Ca^2+^ and firing rate in accordance with coupling strength between neurons [82]. Additionally, natural sensory adaptation also occurs. Consequently, the visual responses observed in the visual cortex reflect the effects of inhibitory/adaptive mechanisms across multiple stages. It is important to note that these mechanisms in the cortex are also activated by ICMS. To achieve a realistic sensation, it may be necessary to utilize these process dynamics in the field of neuroprostheses.

### Hemodynamic changes associated with neural Ca^2+^ signals

In relation to the preceding discussion, quantifying hemodynamic changes in conjunction with induced neural activity provides a better understanding of neurovascular coupling [83] and the contribution of oxygen supply/shortage in neural activity depression. Previous studies have examined hemodynamic changes in the cerebral cortex induced by ICMS [36] and superficial electrical stimulation [84], demonstrating that stronger and longer stimulation leads to greater blood flow changes. Particularly, larger blood flow responses have been reported with optogenetic stimulation of inhibitory neurons [85]. These results are consistent with preferential activation of inhibitory neurons by engaging relatively large blood flow responses that drive large increases in oxyhemoglobin.

Regarding the temporal aspect of the hemodynamic signals, they exhibited notably longer time courses compared to local neural dynamics. During and after the ICMS train, the oxyhemoglobin (reflecting metabolic supply) increased, while the deoxyhemoglobin (reflecting metabolic demand) decreased within the same period. The magnitude of the oxyhemoglobin increase was approximately five times larger than the deoxyhemoglobin decrease. Estimating the relative metabolic demand (OEF) revealed a reduction during ICMS with lower frequencies, particularly at 25 and 50 Hz (Fig. 9e–h). This reduction persisted even after the ICMS train, despite the excitatory neurons exhibiting fewer Ca^2+^ signals and requiring less oxygen compared to the stimulation period. These findings suggest the presence of a metabolic over-supply (hyperoxygenation) environment around the stimulation site that lasts for tens of seconds. The existence of this prolonged hyperoxygenation implies that inhibitory neurons work dominantly during the post-stimulation period. In contrast, such a significant hyperoxygenation was not observed in the visual stimulation condition. As we mentioned above, visual stimulation recruits many sub-cortical circuits, and the full-field LED blinking induced balanced activation and depression. Given that, depression in the full-field visual stimulation condition may reflect the reduction of input from the sub-cortical pathway, requiring relatively less inhibitory neural contribution in the local cortical circuit and relatively less post-stimulation hyperoxygenation in the visual cortex. To achieve a biomimetic response in the field of cortical visual prostheses, it may be necessary to design improved ICMS patterns that effectively balance metabolic supply and demand, at least when representing an environmental illumination.

### Visual prosthesis and the ICMS in the visual cortex

We discovered that the ICMS trains elicited sustained activation, whereas the visual stimulation resulted in only transient responses (e.g., Figure 2). These findings raise questions about the underlying reasons for such stark differences in the induced responses. Moreover, the observed discrepancy in response patterns between ICMS and visual stimulation poses potential challenges in the practical implementation of visual prostheses.

The neurons in the mouse visual cortex exhibited a preference for frequencies around 3 Hz or lower [86, 87]. Thus, the 10-Hz visual stimulation utilized in our study exceeded their neural preferences and could be considered as constant-on visual stimulation for such neurons. Considering the temporal receptive field structures that demonstrate transient on, off, and on/off properties, it becomes evident that the 10-Hz visual stimulation train cannot elicit continuous visual responses following each light flickering. Instead, it generates on and off responses at the onset and offset of the stimulation train. In contrast, ICMS can override these visual preferences in individual neurons. If the aim is to mimic visual responses in visual prostheses, a better approach would involve generating transient responses by adjusting the parameters of ICMS. The simplest method would be to modify the duration of the stimulation train, limiting it to a short period that ensures the necessary activation of neurons [88]. Another option involves exploring the potentially intertwined nature of frequency dependency and the depression effect. Our findings indicate that higher-frequency ICMS induced transient activations that dropped to the weak sustained level within a few seconds after the onset of stimulation (Figs. 6 and S5). Also, in terms of induced response magnitudes, such higher-frequency ICMS led to a smaller activation/depression balance, which was relatively close to the value for visual stimulation (balance ≈ 1; Figure 11l and m). Further increasing the frequency might enhance this trend. However, the perception associated with weak sustained neural responses remains poorly understood. Future studies should address these aspects by employing animal behavior paradigms or by assessing induced perceptions in human subjects.

Paradoxically, natural vision is capable of perceiving constant light. Hence, it is crucial for the visual prosthesis system to generate a constant neural response. Our findings indicate that most of the ICMS frequencies we tested were able to induce significant neural activation throughout the 10-s stimulation train, albeit with varying magnitudes and sizes depending on the frequency employed (Figs. 6 and S5). Some middle frequencies (100 and 250 Hz) resulted in relatively longer activation compared to the other ICMS frequencies. Regarding the magnitude, it remained relatively constant during the sustained period (2–10 s) for middle to high frequencies (e.g., 500 Hz), despite a decreasing trend in the size of the cortical response. Conversely, in terms of the distance, low-frequency ICMS (e.g., 10 Hz) elicited a response of constant size, although the magnitude exhibited an increasing trend. Consequently, it is possible that higher frequencies with increasing pulse intensity or lower frequencies with decreasing pulse intensity could be effective in the ICMS paradigm to generate the constant neural response.

Furthermore, the utilization of depression in the field of visual prostheses can be considered. While depression may prevent continuous light perception or induce perceptual adaptation through the introduction of a refractory period, it could also aid in generating temporally accurate perceptions by rapidly halting unexpectedly prolonged activation. This, in turn, has the potential to enhance the temporal resolution of prosthetic vision, which is currently relatively slower in contemporary visual prostheses or exhibits a lower critical fusion frequency [89]. By effectively inducing depression through the implementation of a higher-frequency ICMS paradigm (Fig. 11l and m) in prosthetic visual systems (e.g., high-frequency burst ICMS evoking temporally localized activation at each burst phase, like blinking-light perception, leading to high temporal resolution and high critical fusion frequency), we could potentially overcome this temporal limitation.

### Tissue damage induced by the probe insertion

Although this study did not specifically investigate the chronic response properties induced by ICMS, certain results suggest that there were effects of tissue damage caused by probe insertion even during the acute experimental phase (see Figure 10aii), which could impair the normal visual response (see Figure S3). Our recordings were conducted approximately 1–2.5 hours after the probe insertion, a period during which processes such as protein binding, neutrophil recruitment, and monocyte recruitment and differentiation into macrophages occur around the surface of the inserted probe [90]. These cellular events can result in the production of reactive oxygen and nitrogen species, which have the potential to harm neurons, as well as cytokines that promote immune responses [91]. Additionally, the traumatic brain injury resulting from probe insertion can lead to acute local ischemia [92]. While these various factors may attenuate normal sensory responses in the sensory cortices, our study demonstrated that ICMS still processes significant capability to induce neural responses, at least during the acute phase. However, during the chronic phase, which includes days following probe insertion, other processes such as foreign body giant cell formation, fibrotic encapsulation around the probe surface [90], and vasculature remodeling [93] occur. These events can have additional effects on neurons and the probes themselves [94]. Further investigation is necessary to examine the potential chronic alterations in neural Ca^2+^ and hemodynamic signals in response to ICMS and sensory stimulation.

## Conclusion

In this study, we made a significant scientific breakthrough by demonstrating that excitatory neurons in the mouse visual cortex exhibit both activation and subsequent depression responses to ICMS. These responses were found to be influenced by the frequencies and modalities of stimulation. Additionally, our analysis of hemodynamic signals revealed that their time courses extended beyond the period of neural depression following activation, exhibiting a large increase in oxygenation that is consistent with the preferential activation of inhibitory neurons and likely suppression of excitatory neural activity. The findings from this work advance our understanding of the fundamental neurophysiology underlying ICMS and provide valuable insights for the development of future neuroprosthetic systems. By uncovering the presence of depression responses and their impact on hemodynamic signals, we open new possibilities for leveraging these phenomena to improve the design and functionality of neuroprosthetic devices. This knowledge has the potential to revolutionize the field of neuroprosthetics and contribute to more effective interventions for individuals with neurological disorders.

## Acknowledgments

The authors would like to thank Christopher L. Hughes, Vanshika Singh, Fan Li, Kevin Stiger, and Camila Garcia for their analysis advice and feedback on this manuscript. This work was supported by NIH R01NS105691, NIH R01NS115707, NIH R01NS129632, NIH R03AG072218, and NSF CAREER CBET 1943906.

## References

[1] N. J. Michelson et al., “Multi-scale, multi-modal analysis uncovers complex relationship at the brain tissue-implant neural interface: new emphasis on the biological interface,” J Neural Eng, vol. 15, no. 3, p. 033001, Jun 2018, doi: 10.1088/1741-2552/aa9dae.

[2] K. Sahasrabuddhe et al., “The Argo: a high channel count recording system for neural recording in vivo,” J Neural Eng, vol. 18, no. 1, p. 015002, Feb 24 2021, doi: 10.1088/1741-2552/abd0ce.

[3] N. Suematsu, T. Naito, T. Miyoshi, H. Sawai, and H. Sato, “Spatiotemporal receptive field structures in retinogeniculate connections of cat,” Front Syst Neurosci, vol. 7, p. 103, 2013, doi: 10.3389/fnsys.2013.00103.

[4] N. Suematsu, T. Naito, and H. Sato, “Relationship between orientation sensitivity and spatiotemporal receptive field structures of neurons in the cat lateral geniculate nucleus,” Neural Netw, vol. 35, pp. 10–20, Nov 2012, doi: 10.1016/j.neunet.2012.06.008.

[5] M. H. Histed, V. Bonin, and R. C. Reid, “Direct activation of sparse, distributed populations of cortical neurons by electrical microstimulation,” Neuron, vol. 63, no. 4, pp. 508–22, Aug 27 2009, doi: 10.1016/j.neuron.2009.07.016.

[6] S. Miyamoto, N. Suematsu, Y. Umehira, Y. Hayashida, and T. Yagi, “Age-related changes in the spatiotemporal responses to electrical stimulation in the visual cortex of rats with progressive vision loss,” Sci Rep, vol. 7, no. 1, p. 14165, Oct 26 2017, doi: 10.1038/s41598-017-14303-1.

[7] S. D. Stoney, Jr., W. D. Thompson, and H. Asanuma, “Excitation of pyramidal tract cells by intracortical microstimulation: effective extent of stimulating current,” J Neurophysiol, vol. 31, no. 5, pp. 659–69, Sep 1968, doi: 10.1152/jn.1968.31.5.659.

[8] E. J. Tehovnik, A. S. Tolias, F. Sultan, W. M. Slocum, and N. K. Logothetis, “Direct and indirect activation of cortical neurons by electrical microstimulation,” J Neurophysiol, vol. 96, no. 2, pp. 512–21, Aug 2006, doi: 10.1152/jn.00126.2006.

[9] S. N. Flesher et al., “Intracortical microstimulation of human somatosensory cortex,” Sci Transl Med, vol. 8, no. 361, p. 361ra141, Oct 19 2016, doi: 10.1126/scitranslmed.aaf8083.

[10] C. L. Hughes, S. N. Flesher, and R. A. Gaunt, “Effects of stimulus pulse rate on somatosensory adaptation in the human cortex,” Brain Stimul, vol. 15, no. 4, pp. 987–995, Jul-Aug 2022, doi: 10.1016/j.brs.2022.05.021.

[11] C. L. Hughes et al., “Neural stimulation and recording performance in human sensorimotor cortex over 1500 days,” J Neural Eng, vol. 18, no. 4, Aug 13 2021, doi: 10.1088/1741-2552/ac18ad.

[12] T. Allison-Walker, M. A. Hagan, N. S. C. Price, and Y. T. Wong, “Microstimulation-evoked neural responses in visual cortex are depth dependent,” Brain Stimul, vol. 14, no. 4, pp. 741–750, Jul-Aug 2021, doi: 10.1016/j.brs.2021.04.020.

[13] S. J. Meikle, M. Ann Hagan, N. S. Chiang Price, and Y. Tat Wong, “Filling in the Visual Gaps: Shifting Cortical Activity using Current Steering,” Annu Int Conf IEEE Eng Med Biol Soc, vol. 2021, pp. 5733–5736, Nov 2021, doi: 10.1109/EMBC46164.2021.9629693.

[14] S. J. Meikle, M. A. Hagan, N. S. C. Price, and Y. T. Wong, “Intracortical current steering shifts the location of evoked neural activity,” J Neural Eng, vol. 19, no. 3, Jun 23 2022, doi: 10.1088/1741-2552/ac77bf.

[15] X. Chen, F. Wang, E. Fernandez, and P. R. Roelfsema, “Shape perception via a high-channel-count neuroprosthesis in monkey visual cortex,” Science, vol. 370, no. 6521, pp. 1191–1196, Dec 4 2020, doi: 10.1126/science.abd7435.

[16] T. S. Davis et al., “Spatial and temporal characteristics of V1 microstimulation during chronic implantation of a microelectrode array in a behaving macaque,” J Neural Eng, vol. 9, no. 6, p. 065003, Dec 2012, doi: 10.1088/1741-2560/9/6/065003.

[17] P. H. Schiller, W. M. Slocum, M. C. Kwak, G. L. Kendall, and E. J. Tehovnik, “New methods devised specify the size and color of the spots monkeys see when striate cortex (area V1) is electrically stimulated,” Proc Natl Acad Sci U S A, vol. 108, no. 43, pp. 17809–14, Oct 25 2011, doi: 10.1073/pnas.1108337108.

[18] K. Torab, T. S. Davis, D. J. Warren, P. A. House, R. A. Normann, and B. Greger, “Multiple factors may influence the performance of a visual prosthesis based on intracortical microstimulation: nonhuman primate behavioural experimentation,” J Neural Eng, vol. 8, no. 3, p. 035001, Jun 2011, doi: 10.1088/1741-2560/8/3/035001.

[19] M. Armenta Salas et al., “Sequence of visual cortex stimulation affects phosphene brightness in blind subjects,” Brain Stimul, vol. 15, no. 3, pp. 605–614, May-Jun 2022, doi: 10.1016/j.brs.2022.03.008.

[20] E. Fernandez et al., “Visual percepts evoked with an intracortical 96-channel microelectrode array inserted in human occipital cortex,” J Clin Invest, vol. 131, no. 23, Dec 1 2021, doi: 10.1172/JCI151331.

[21] A. J. Lowery et al., “Monash Vision Group’s Gennaris Cortical Implant for Vision Restoration,” in Artificial Vision: A Clinical Guide, V. P. Gabel Ed. Cham: Springer International Publishing, 2017, pp. 215–225.

[22] E. M. Schmidt, M. J. Bak, F. T. Hambrecht, C. V. Kufta, D. K. O’Rourke, and P. Vallabhanath, “Feasibility of a visual prosthesis for the blind based on intracortical microstimulation of the visual cortex,” Brain, vol. 119 ( Pt 2), pp. 507–22, Apr 1996, doi: 10.1093/brain/119.2.507.

[23] P. R. Troyk, “The Intracortical Visual Prosthesis Project,” in Artificial Vision: A Clinical Guide, V. P. Gabel Ed. Cham: Springer International Publishing, 2017, pp. 203–214.

[24] J. R. Eles and T. D. Y. Kozai, “In vivo imaging of calcium and glutamate responses to intracortical microstimulation reveals distinct temporal responses of the neuropil and somatic compartments in layer II/III neurons,” Biomaterials, vol. 234, p. 119767, Mar 2020, doi: 10.1016/j.biomaterials.2020.119767.

[25] J. R. Eles, K. C. Stieger, and T. D. Y. Kozai, “The temporal pattern of intracortical microstimulation pulses elicits distinct temporal and spatial recruitment of cortical neuropil and neurons,” J Neural Eng, vol. 18, no. 1, Jan 25 2021, doi: 10.1088/1741-2552/abc29c.

[26] Z. Ma et al., “Two-photon calcium imaging of neuronal and astrocytic responses: the influence of electrical stimulus parameters and calcium signaling mechanisms,” J Neural Eng, vol. 18, no. 4, Jul 2 2021, doi: 10.1088/1741-2552/ac0b50.

[27] N. J. Michelson, J. R. Eles, A. L. Vazquez, K. A. Ludwig, and T. D. Y. Kozai, “Calcium activation of cortical neurons by continuous electrical stimulation: Frequency dependence, temporal fidelity, and activation density,” J Neurosci Res, vol. 97, no. 5, pp. 620–638, May 2019, doi: 10.1002/jnr.24370.

[28] K. C. Stieger, J. R. Eles, K. A. Ludwig, and T. D. Y. Kozai, “In vivo microstimulation with cathodic and anodic asymmetric waveforms modulates spatiotemporal calcium dynamics in cortical neuropil and pyramidal neurons of male mice,” J Neurosci Res, vol. 98, no. 10, pp. 2072–2095, Oct 2020, doi: 10.1002/jnr.24676.

[29] K. C. Stieger, J. R. Eles, K. A. Ludwig, and T. D. Y. Kozai, “Intracortical microstimulation pulse waveform and frequency recruits distinct spatiotemporal patterns of cortical neuron and neuropil activation,” J Neural Eng, vol. 19, no. 2, Mar 31 2022, doi: 10.1088/1741-2552/ac5bf5.

[30] K. Kumaravelu, J. Sombeck, L. E. Miller, S. J. Bensmaia, and W. M. Grill, “Stoney vs. Histed: Quantifying the spatial effects of intracortical microstimulation,” Brain Stimul, vol. 15, no. 1, pp. 141–151, Jan-Feb 2022, doi: 10.1016/j.brs.2021.11.015.

[31] S. Butovas, S. G. Hormuzdi, H. Monyer, and C. Schwarz, “Effects of electrically coupled inhibitory networks on local neuronal responses to intracortical microstimulation,” J Neurophysiol, vol. 96, no. 3, pp. 1227–36, Sep 2006, doi: 10.1152/jn.01170.2005.

[32] D. B. McCreery, T. G. Yuen, W. F. Agnew, and L. A. Bullara, “A characterization of the effects on neuronal excitability due to prolonged microstimulation with chronically implanted microelectrodes,” IEEE Trans Biomed Eng, vol. 44, no. 10, pp. 931–9, Oct 1997, doi: 10.1109/10.634645.

[33] G. K. Wu, Y. Ardeshirpour, C. Mastracchio, J. Kent, M. Caiola, and M. Ye, “Amplitude– and frequency-dependent activation of layer II/III neurons by intracortical microstimulation,” iScience, vol. 26, no. 11, p. 108140, Nov 17 2023, doi: 10.1016/j.isci.2023.108140.

[34] C. D. Eiber, N. H. Lovell, and G. J. Suaning, “Attaining higher resolution visual prosthetics: a review of the factors and limitations,” J Neural Eng, vol. 10, no. 1, p. 011002, Feb 2013, doi: 10.1088/1741-2560/10/1/011002.

[35] Y. Ma et al., “Wide-field optical mapping of neural activity and brain haemodynamics: considerations and novel approaches,” Philos Trans R Soc Lond B Biol Sci, vol. 371, no. 1705, Oct 5 2016, doi: 10.1098/rstb.2015.0360.

[36] A. A. Brock, R. M. Friedman, R. H. Fan, and A. W. Roe, “Optical imaging of cortical networks via intracortical microstimulation,” J Neurophysiol, vol. 110, no. 11, pp. 2670–8, Dec 2013, doi: 10.1152/jn.00879.2012.

[37] H. Dana et al., “Thy1-GCaMP6 transgenic mice for neuronal population imaging in vivo,” PLoS One, vol. 9, no. 9, p. e108697, 2014, doi: 10.1371/journal.pone.0108697.

[38] S. Kura, H. Xie, B. Fu, C. Ayata, D. A. Boas, and S. Sakadzic, “Intrinsic optical signal imaging of the blood volume changes is sufficient for mapping the resting state functional connectivity in the rodent cortex,” J Neural Eng, vol. 15, no. 3, p. 035003, Jun 2018, doi: 10.1088/1741-2552/aaafe4.

[39] A. Edelstein, N. Amodaj, K. Hoover, R. Vale, and N. Stuurman, “Computer control of microscopes using microManager,” Curr Protoc Mol Biol, vol. Chapter 14, p. Unit14 20, Oct 2010, doi: 10.1002/0471142727.mb1420s92.

[40] N. Stuurman, N. Amdodaj, and R. Vale, “μManager: Open Source Software for Light Microscope Imaging,” Microscopy Today, vol. 15, no. 3, pp. 42–43, 2007, doi: 10.1017/s1551929500055541.

[41] D. R. Merrill, M. Bikson, and J. G. Jefferys, “Electrical stimulation of excitable tissue: design of efficacious and safe protocols,” J Neurosci Methods, vol. 141, no. 2, pp. 171–98, Feb 15 2005, doi: 10.1016/j.jneumeth.2004.10.020.

[42] J. Schindelin et al., “Fiji: an open-source platform for biological-image analysis,” Nat Methods, vol. 9, no. 7, pp. 676–82, Jun 28 2012, doi: 10.1038/nmeth.2019.

[43] J. Canny, “A Computational Approach to Edge Detection,” IEEE Transactions on Pattern Analysis and Machine Intelligence, vol. PAMI-8, no. 6, pp. 679–698, 1986, doi: 10.1109/tpami.1986.4767851.

[44] Q. Wang, D. C. Millard, H. J. Zheng, and G. B. Stanley, “Voltage-sensitive dye imaging reveals improved topographic activation of cortex in response to manipulation of thalamic microstimulation parameters,” J Neural Eng, vol. 9, no. 2, p. 026008, Apr 2012, doi: 10.1088/1741-2560/9/2/026008.

[45] D. Attwell and C. Iadecola, “The neural basis of functional brain imaging signals,” Trends Neurosci, vol. 25, no. 12, pp. 621–5, Dec 2002, doi: 10.1016/s0166-2236(02)02264-6.

[46] D. B. McCreery, L. A. Bullara, and W. F. Agnew, “Neuronal activity evoked by chronically implanted intracortical microelectrodes,” Exp Neurol, vol. 92, no. 1, pp. 147–61, Apr 1986, doi: 10.1016/0014-4886(86)90131-7.

[47] D. B. McCreery, T. G. Yuen, W. F. Agnew, and L. A. Bullara, “Stimulation with chronically implanted microelectrodes in the cochlear nucleus of the cat: histologic and physiologic effects,” Hear Res, vol. 62, no. 1, pp. 42–56, Sep 1992, doi: 10.1016/0378-5955(92)90201-w.

[48] D. B. McCreery and W. F. Agnew, “Changes in extracellular potassium and calcium concentration and neural activity during prolonged electrical stimulation of the cat cerebral cortex at defined charge densities,” Exp Neurol, vol. 79, no. 2, pp. 371–96, Feb 1983, doi: 10.1016/0014-4886(83)90220-0.

[49] A. A. P. Leao, “Spreading Depression of Activity in the Cerebral Cortex,” Journal of Neurophysiology, vol. 7, no. 6, pp. 359–390, 1944, doi: 10.1152/jn.1944.7.6.359.

[50] R. S. McLachlan and J. P. Girvin, “Spreading depression of Leao in rodent and human cortex,” Brain Res, vol. 666, no. 1, pp. 133–6, Dec 12 1994, doi: 10.1016/0006-8993(94)90295-x.

[51] A. M. Ba et al., “Multiwavelength optical intrinsic signal imaging of cortical spreading depression,” J Neurophysiol, vol. 88, no. 5, pp. 2726–35, Nov 2002, doi: 10.1152/jn.00729.2001.

[52] A. de Crespigny, J. Rother, N. van Bruggen, C. Beaulieu, and M. E. Moseley, “Magnetic resonance imaging assessment of cerebral hemodynamics during spreading depression in rats,” J Cereb Blood Flow Metab, vol. 18, no. 9, pp. 1008–17, Sep 1998, doi: 10.1097/00004647-199809000-00010.

[53] T. Houben et al., “Optogenetic induction of cortical spreading depression in anesthetized and freely behaving mice,” J Cereb Blood Flow Metab, vol. 37, no. 5, pp. 1641–1655, May 2017, doi: 10.1177/0271678X16645113.

[54] A. Van Harreveld and S. Ochs, “Electrical and vascular concomitants of spreading depression,” Am J Physiol, vol. 189, no. 1, pp. 159–66, Apr 1957, doi: 10.1152/ajplegacy.1957.189.1.159.

[55] P. M. Sawant-Pokam, P. Suryavanshi, J. M. Mendez, F. E. Dudek, and K. C. Brennan, “Mechanisms of Neuronal Silencing After Cortical Spreading Depression,” Cereb Cortex, vol. 27, no. 2, pp. 1311–1325, Feb 1 2017, doi: 10.1093/cercor/bhv328.

[56] R. Yun, J. H. Mishler, S. I. Perlmutter, R. P. N. Rao, and E. E. Fetz, “Responses of cortical neurons to intracortical microstimulation in awake primates,” bioRxiv, 2022, doi: 10.1101/2022.03.30.486457.

[57] H. Markram, M. Toledo-Rodriguez, Y. Wang, A. Gupta, G. Silberberg, and C. Wu, “Interneurons of the neocortical inhibitory system,” Nat Rev Neurosci, vol. 5, no. 10, pp. 793–807, Oct 2004, doi: 10.1038/nrn1519.

[58] R. Tremblay, S. Lee, and B. Rudy, “GABAergic Interneurons in the Neocortex: From Cellular Properties to Circuits,” Neuron, vol. 91, no. 2, pp. 260–92, Jul 20 2016, doi: 10.1016/j.neuron.2016.06.033.

[59] M. Matos et al., “Astrocytes detect and upregulate transmission at inhibitory synapses of somatostatin interneurons onto pyramidal cells,” Nat Commun, vol. 9, no. 1, p. 4254, Oct 12 2018, doi: 10.1038/s41467-018-06731-y.

[60] J. Lee et al., “Opposed hemodynamic responses following increased excitation and parvalbumin-based inhibition,” J Cereb Blood Flow Metab, vol. 41, no. 4, pp. 841–856, Apr 2021, doi: 10.1177/0271678X20930831.

[61] C. Neudorfer et al., “Kilohertz-frequency stimulation of the nervous system: A review of underlying mechanisms,” Brain Stimul, vol. 14, no. 3, pp. 513–530, May-Jun 2021, doi: 10.1016/j.brs.2021.03.008.

[62] J. K. Thompson, M. R. Peterson, and R. D. Freeman, “Single-neuron activity and tissue oxygenation in the cerebral cortex,” Science, vol. 299, no. 5609, pp. 1070–2, Feb 14 2003, doi: 10.1126/science.1079220.

[63] T. Abrahamsson, B. Gustafsson, and E. Hanse, “Synaptic fatigue at the naive perforant path-dentate granule cell synapse in the rat,” J Physiol, vol. 569, no. Pt 3, pp. 737–50, Dec 15 2005, doi: 10.1113/jphysiol.2005.097725.

[64] J. A. Gomez et al., “Ventral tegmental area astrocytes orchestrate avoidance and approach behavior,” Nat Commun, vol. 10, no. 1, p. 1455, Mar 29 2019, doi: 10.1038/s41467-019-09131-y.

[65] Y. Tanaka et al., “Focal activation of neuronal circuits induced by microstimulation in the visual cortex,” J Neural Eng, vol. 16, no. 3, p. 036007, Jun 2019, doi: 10.1088/1741-2552/ab0b80.

[66] M. Avermann, C. Tomm, C. Mateo, W. Gerstner, and C. C. Petersen, “Microcircuits of excitatory and inhibitory neurons in layer 2/3 of mouse barrel cortex,” J Neurophysiol, vol. 107, no. 11, pp. 3116–34, Jun 2012, doi: 10.1152/jn.00917.2011.

[67] A. Karagiannis et al., “Lactate is an energy substrate for rodent cortical neurons and enhances their firing activity,” Elife, vol. 10, Nov 12 2021, doi: 10.7554/eLife.71424.

[68] S. Mason, “Lactate Shuttles in Neuroenergetics-Homeostasis, Allostasis and Beyond,” Front Neurosci, vol. 11, p. 43, 2017, doi: 10.3389/fnins.2017.00043.

[69] M. Tsacopoulos and P. J. Magistretti, “Metabolic coupling between glia and neurons,” J Neurosci, vol. 16, no. 3, pp. 877–85, Feb 1 1996, doi: 10.1523/JNEUROSCI.16-03-00877.1996.

[70] L. D. Pozzo-Miller et al., “Impairments in high-frequency transmission, synaptic vesicle docking, and synaptic protein distribution in the hippocampus of BDNF knockout mice,” J Neurosci, vol. 19, no. 12, pp. 4972–83, Jun 15 1999, doi: 10.1523/JNEUROSCI.19-12-04972.1999.

[71] J. E. Cohen and R. D. Fields, “Extracellular calcium depletion in synaptic transmission,” Neuroscientist, vol. 10, no. 1, pp. 12–7, Feb 2004, doi: 10.1177/1073858403259440.

[72] S. C. Bellinger, G. Miyazawa, and P. N. Steinmetz, “Submyelin potassium accumulation may functionally block subsets of local axons during deep brain stimulation: a modeling study,” J Neural Eng, vol. 5, no. 3, pp. 263–74, Sep 2008, doi: 10.1088/1741-2560/5/3/001.

[73] T. V. Bliss and S. F. Cooke, “Long-term potentiation and long-term depression: a clinical perspective,” Clinics (Sao Paulo), vol. 66 Suppl 1, no. Suppl 1, pp. 3–17, 2011, doi: 10.1590/s1807-59322011001300002.

[74] C. Mateo, P. M. Knutsen, P. S. Tsai, A. Y. Shih, and D. Kleinfeld, “Entrainment of Arteriole Vasomotor Fluctuations by Neural Activity Is a Basis of Blood-Oxygenation-Level-Dependent “Resting-State” Connectivity,” Neuron, vol. 96, no. 4, pp. 936–948 e3, Nov 15 2017, doi: 10.1016/j.neuron.2017.10.012.

[75] P. J. O’Herron, D. A. Hartmann, K. Xie, P. Kara, and A. Y. Shih, “3D optogenetic control of arteriole diameter in vivo,” Elife, vol. 11, Sep 15 2022, doi: 10.7554/eLife.72802.

[76] T. J. Richner et al., “Patterned optogenetic modulation of neurovascular and metabolic signals,” J Cereb Blood Flow Metab, vol. 35, no. 1, pp. 140–7, Jan 2015, doi: 10.1038/jcbfm.2014.189.

[77] J. S. Marvin et al., “Stability, affinity, and chromatic variants of the glutamate sensor iGluSnFR,” Nat Methods, vol. 15, no. 11, pp. 936–939, Nov 2018, doi: 10.1038/s41592-018-0171-3.

[78] J. S. Marvin et al., “A genetically encoded fluorescent sensor for in vivo imaging of GABA,” Nat Methods, vol. 16, no. 8, pp. 763–770, Aug 2019, doi: 10.1038/s41592-019-0471-2.

[79] J. Yeonan-Kim and M. Bertalmio, “Retinal Lateral Inhibition Provides the Biological Basis of Long-Range Spatial Induction,” PLoS One, vol. 11, no. 12, p. e0168963, 2016, doi: 10.1371/journal.pone.0168963.

[80] J. M. Budd and Z. F. Kisvarday, “Local lateral connectivity of inhibitory clutch cells in layer 4 of cat visual cortex (area 17),” Exp Brain Res, vol. 140, no. 2, pp. 245–50, Sep 2001, doi: 10.1007/s002210100817.

[81] W. Singer and O. D. Creutzfeldt, “Reciprocal lateral inhibition of on– and off-center neurones in the lateral geniculate body of the cat,” Exp Brain Res, vol. 10, no. 3, pp. 311–30, 1970, doi: 10.1007/BF00235054.

[82] J. F. O’Rawe, Z. Zhou, A. J. Li, P. K. LaFosse, H. C. Goldbach, and M. H. Histed, “Excitation creates a distributed pattern of cortical suppression due to varied recurrent input,” Neuron, Oct 20 2023, doi: 10.1016/j.neuron.2023.09.010.

[83] J. A. Padawer-Curry et al., “Psychedelic 5-HT2A receptor agonism: neuronal signatures and altered neurovascular coupling,” bioRxiv, 2023, doi: 10.1101/2023.09.23.559145.

[84] M. Suh, S. Bahar, A. D. Mehta, and T. H. Schwartz, “Blood volume and hemoglobin oxygenation response following electrical stimulation of human cortex,” Neuroimage, vol. 31, no. 1, pp. 66–75, May 15 2006, doi: 10.1016/j.neuroimage.2005.11.030.

[85] A. L. Vazquez, M. Fukuda, and S. G. Kim, “Inhibitory Neuron Activity Contributions to Hemodynamic Responses and Metabolic Load Examined Using an Inhibitory Optogenetic Mouse Model,” Cereb Cortex, vol. 28, no. 11, pp. 4105–4119, Nov 1 2018, doi: 10.1093/cercor/bhy225.

[86] D. Camillo, M. Ahmadlou, and J. A. Heimel, “Contrast-Dependence of Temporal Frequency Tuning in Mouse V1,” Front Neurosci, vol. 14, p. 868, 2020, doi: 10.3389/fnins.2020.00868.

[87] H. Wang, O. Dey, W. N. Lagos, and E. M. Callaway, “Diversity in spatial frequency, temporal frequency, and speed tuning across mouse visual cortical areas and layers,” J Comp Neurol, vol. 530, no. 18, pp. 3226–3247, Dec 2022, doi: 10.1002/cne.25404.

[88] C. Hughes and T. Kozai, “Dynamic amplitude modulation of microstimulation evokes biomimetic onset and offset transients and reduces depression of evoked calcium responses in sensory cortices,” Brain Stimul, vol. 16, no. 3, pp. 939–965, May-Jun 2023, doi: 10.1016/j.brs.2023.05.013.

[89] E. Borda and D. Ghezzi, “Advances in visual prostheses: engineering and biological challenges,” Progress in Biomedical Engineering, vol. 4, no. 3, 2022, doi: 10.1088/2516-1091/ac812c.

[90] A. Carnicer-Lombarte, S. T. Chen, G. G. Malliaras, and D. G. Barone, “Foreign Body Reaction to Implanted Biomaterials and Its Impact in Nerve Neuroprosthetics,” Front Bioeng Biotechnol, vol. 9, p. 622524, 2021, doi: 10.3389/fbioe.2021.622524.

[91] A. Kanashiro et al., “The role of neutrophils in neuro-immune modulation,” Pharmacol Res, vol. 151, p. 104580, Jan 2020, doi: 10.1016/j.phrs.2019.104580.

[92] T. D. Kozai et al., “Reduction of neurovascular damage resulting from microelectrode insertion into the cerebral cortex using in vivo two-photon mapping,” J Neural Eng, vol. 7, no. 4, p. 046011, Aug 2010, doi: 10.1088/1741-2560/7/4/046011.

[93] A. Salehi, J. H. Zhang, and A. Obenaus, “Response of the cerebral vasculature following traumatic brain injury,” J Cereb Blood Flow Metab, vol. 37, no. 7, pp. 2320–2339, Jul 2017, doi: 10.1177/0271678X17701460.

[94] T. D. Kozai et al., “Mechanical failure modes of chronically implanted planar silicon-based neural probes for laminar recording,” Biomaterials, vol. 37, pp. 25–39, Jan 2015, doi: 10.1016/j.biomaterials.2014.10.040.

