## Supplemental Data for "Activation and depression of neural and hemodynamic responses induced by the intracortical microstimulation and visual stimulation in the mouse visual cortex"

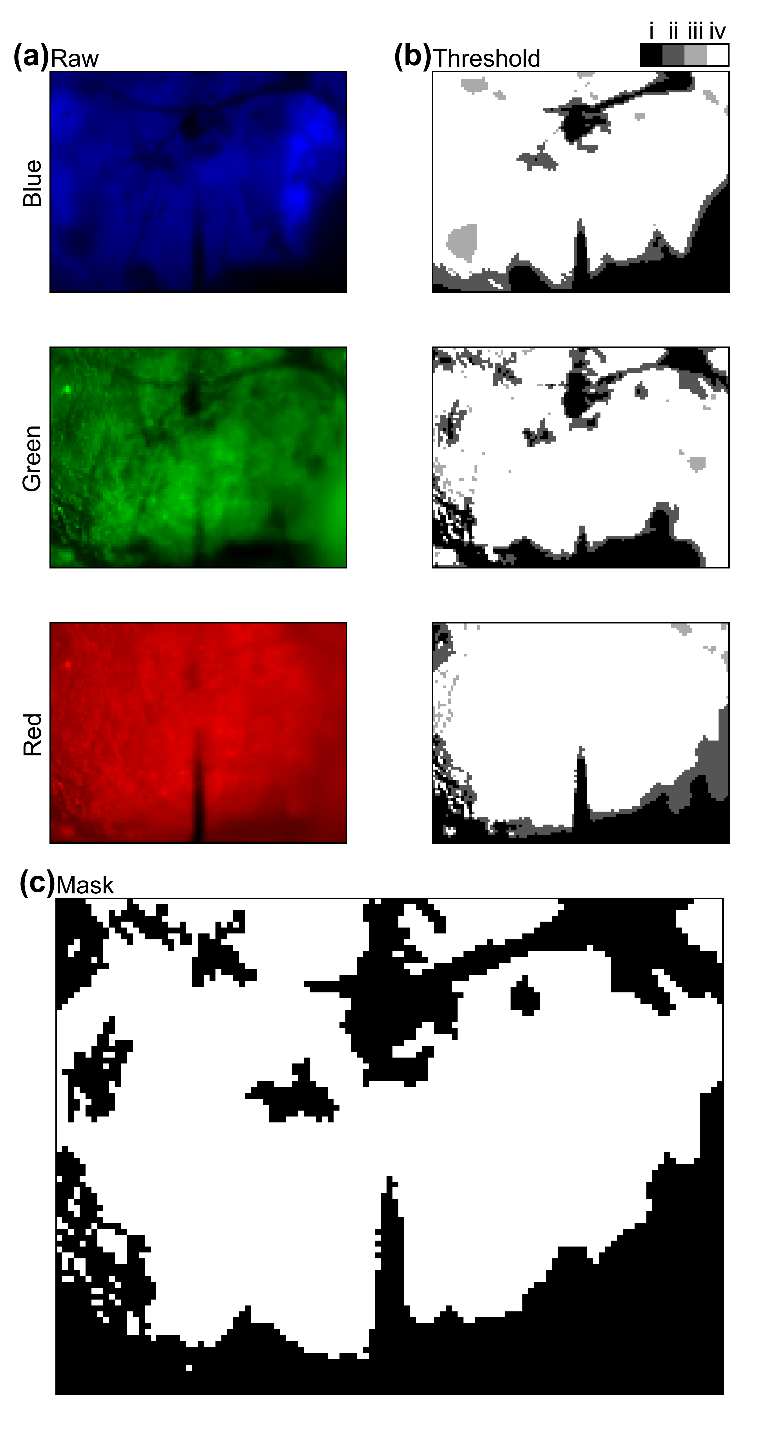

**Figure S1. Vessel pattern masking.**

(a) averaged raw images taken under blue-, green-, and red-light illuminations. (b) hysteresis thresholding with two different intensity levels for effective extraction of connecting regions. (i) regions darker than mean-1SD, (ii) regions darker than mean-0.5SD, brighter than mean-1SD, connecting with (i), (iii) regions darker than mean-0.5SD, brighter than mean-1SD, separating from (i), (iv) regions brighter than mean-0.5SD. Regions (i) and (ii) are defined as vessel patterns and poorly illuminated locations like probe shank and edges of an image, and excluded from further analyses.

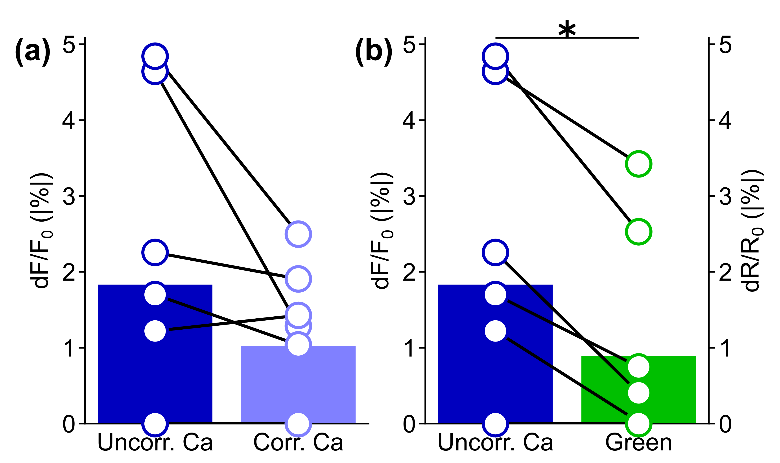

**Figure S2. Hemodynamic correction of Ca^2+^ signals**

(a) Comparison of depression magnitudes between uncorrected and corrected Ca^2+^ signals (paired t-test, *t* = 1.75, *p* = 0.12). (b) Comparison of depression magnitudes between uncorrected Ca^2+^ and green signals (*t* = 3.00, *p* < 0.05). Depression period and area are derived from the results of corrected Ca^2+^. Uncorrected and corrected Ca^2+^ signals are shown in fluorescent change ratio (dF/F_0_). Green signal is shown in reflectance change ratio (dR/R_0_). Although depression magnitude in corrected Ca^2+^ signal decreased compared to uncorrected Ca^2+^ signal, it is still significantly greater than zero (one-sample t-test, *t* = 3.03, *p* < 0.05).

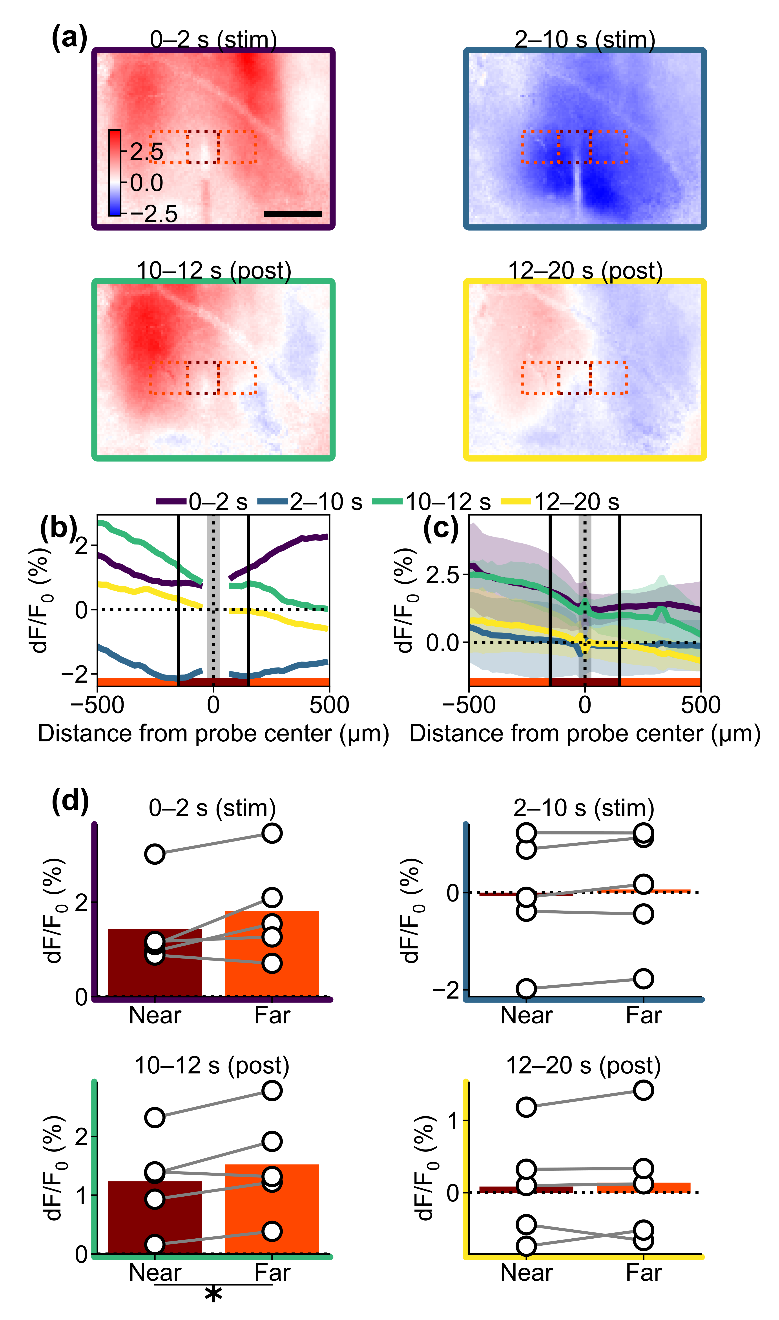

**Figure S3: Visually-evoked response was suppressed around the inserted probes.**

(a) Representative examples of XY-response map at the transient onset (0–2 s, during stimulation), sustained onset (2–10 s, during stimulation), transient offset (10–12, post-stimulation), and sustained offset (12–20 s, post-stimulation) periods of the visual stimulation train. (b) Line profiles of the response around the probe (averaged over 300 µm along the shank, vertical axis of dotted-line boxes in a). Vertical solid lines indicate boundaries dividing “near” (27.5–150 µm) and “far” (150–500 µm) locations from the stimulation channel center. Gray shaded area represents the inserted probe shank (±27.5 µm). Line colors correspond to the time points after the stimulation onset, shown in (a). (c) Population line profile (mean ± 1SD among the animals, n = 5). (d) Comparisons of the visual response magnitudes between the near and far locations from the probe shank center. One-tailed paired t-test revealed significant response reductions in the near region at transient offset (10–12 s, *t* = -2.66, *p* < 0.05) periods. * indicates *p* < 0.05.

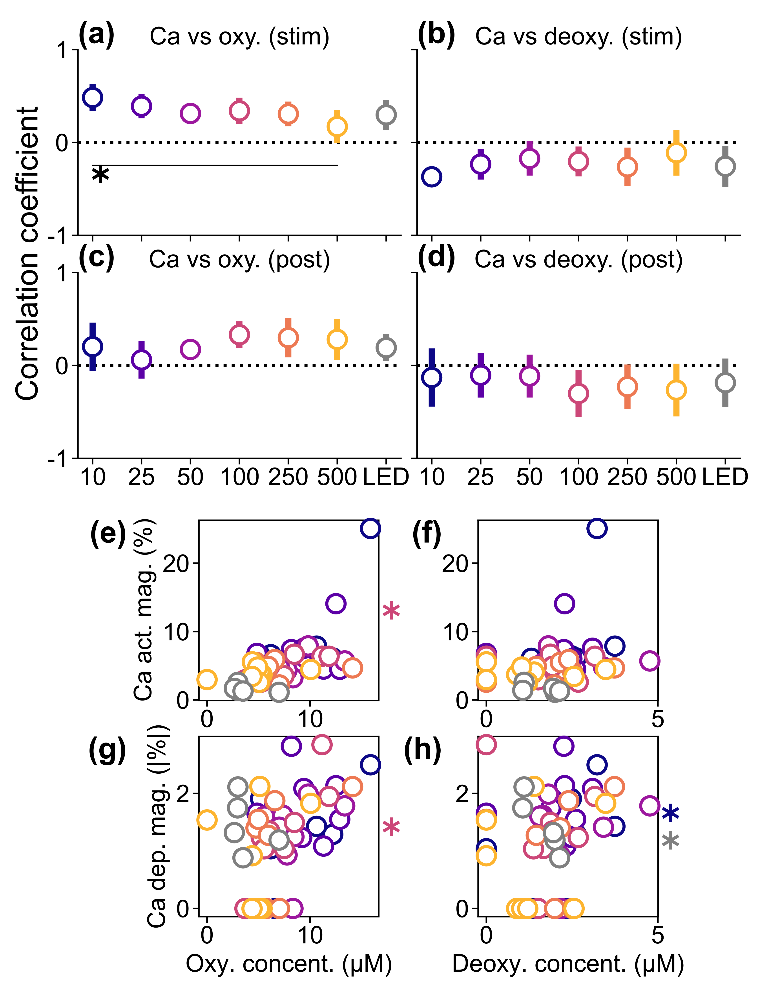

**Figure S4. Ca^2+^ and hemodynamic spatiotemporal correlation.**

(a) Averaged Spearman’s correlation coefficients (mean ± 1SD among animals) between Ca^2+^ and oxyhemoglobin spatiotemporal signals during the stimulation period (0–10 s from the stimulation onset), which was significantly affected by the stimulation conditions (one-way ANOVA, *F* = 3.02, *p* < 0.05). Post-hoc Tukey’s HSD test revealed a significant difference between 10-Hz ICMS and 500-Hz ICMS conditions (difference = -0.31, *p* < 0.05). (b) Averaged Spearman’s correlation coefficients between Ca^2+^ and deoxyhemoglobin spatiotemporal signals during the stimulation period, which were not significantly affected by the stimulation conditions (*F* = 1.29, *p* = 0.28). (c) Averaged Spearman’s correlation coefficients between Ca^2+^ and oxyhemoglobin spatiotemporal signals during the post-stimulation period (10–20 s from the stimulation onset), which was not significantly affected by the stimulation conditions (*F* = 1.61, *p* = 0.17). (d) Averaged Spearman’s correlation coefficients between Ca^2+^ and deoxyhemoglobin spatiotemporal signals during the post-stimulation period, which were not significantly affected by the stimulation conditions (*F* = 0.61, *p* = 0.72). (e) Comparisons between the Ca^2+^ activation magnitude and the oxyhemoglobin concentration change in the various stimulation conditions (colors correspond to a–d). Each circle indicates each mouse. (f) Comparisons between the Ca^2+^ activation magnitude and the deoxyhemoglobin concentration change. (g) Comparisons between the Ca^2+^ depression magnitude and the oxyhemoglobin concentration change. (h) Comparisons between the Ca^2+^ depression magnitude and the deoxyhemoglobin concentration change. * with horizontal line indicates *p* < 0.05 for pairwise comparison with post-hoc Tukey’s HSD test. * in e–h indicates significant correlation; colors correspond to stimulation conditions.

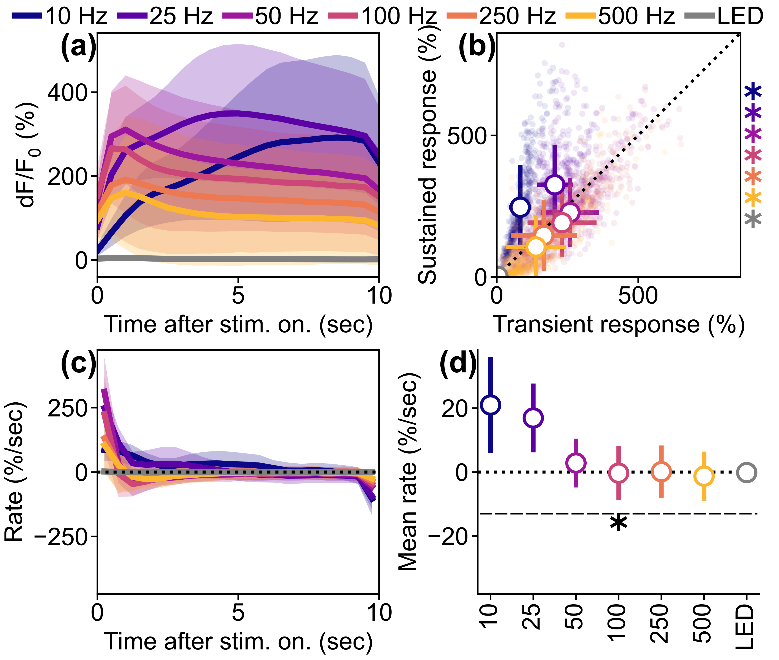

**Figure S5. Temporal dynamics of the induced activations in soma.**

(a) Time courses of the activation in soma under the various stimulation conditions. Lines and shaded areas indicate mean ± 1SD. (b) Comparisons of the response magnitudes between the transient (0–2 s) and the sustained (2–10 s) periods. Dots represent each soma. Circles with error bars indicate mean ± 1SD. The 10- and 25-Hz ICMSs induced significantly stronger sustained responses than transient ones (paired t-test, 10 Hz, *t* = -28.92, *p* < 0.05; 25 Hz, *t* = -21.98, *p* < 0.05). The 50–500-Hz ICMSs and visual stimulation, on the other hand, elicited significantly stronger transient responses than sustained ones (50 Hz, *t* = 12.28, *p* < 0.05; 100 Hz, *t* = 13.62, *p* < 0.05; 250 Hz, *t* = 6.95, *p* < 0.05; 500 Hz, *t* = 10.88, *p* < 0.05; LED, *t* = 7.98, *p* < 0.05). (c) Change rates of the activation magnitude (mean ± 1SD). (d) Mean change rates among the stimulation conditions over the 10-sec stimulation period (mean ± 1SD). Mean change rates of the activation magnitude significantly varied depending on the stimulation conditions (one-way ANOVA, *F* = 395.77, *p* < 0.05), where the 10–50 Hz ICMSs induced significantly higher change rates than the other conditions (post-hoc Tukey’s HSD test, see Table S1 for detail).

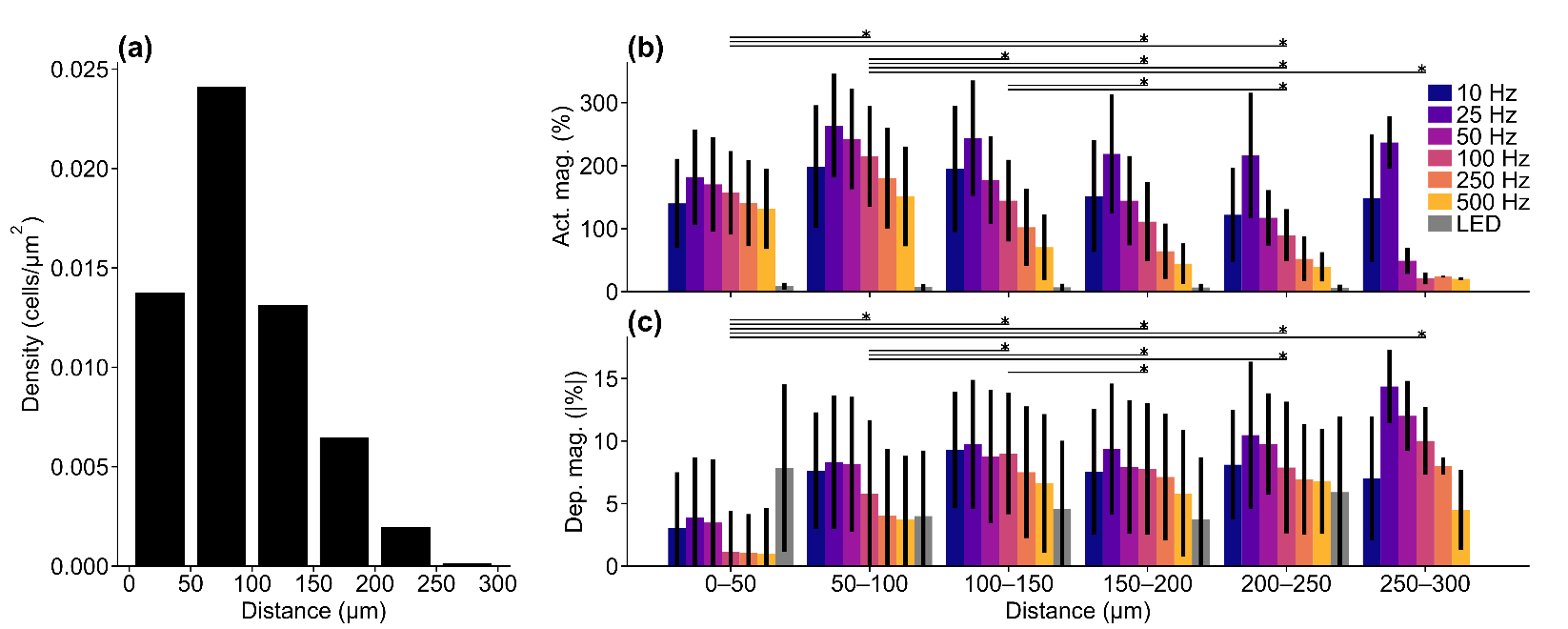

**Figure S6. Distance dependency of the somatic activation and depression.**

(a) Identified ROI density over distance from stimulation channels center to a center of mass of each ROI. (b and c) Activation and depression magnitudes over the distances (mean ± 1SD among cells). Both distance and stimulation condition significantly modulate the activation and depression magnitudes of somatic Ca^2+^ signals (two-way ANOVA, activation, *F*_distance_ = 52.29, *p*_distance_ < 0.05, *F*_stimulation condition_ = 334.34, *p*_stimulation condition_ < 0.05, *F*_distance × stimulation condition_ = 6.01, *p*_distance × stimulation condition_ < 0.05; depression, *F*_distance_ = 5.14, *p*_distance_ < 0.05, *F*_stimulation condition_ = 17.62, *p*_stimulation condition_ < 0.05, *F*_distance × stimulation condition_ = 2.24, *p*_distance × stimulation condition_ < 0.05). * with horizontal line indicates *p* < 0.05 for pairwise comparison between distances with post-hoc Tukey’s HSD test.

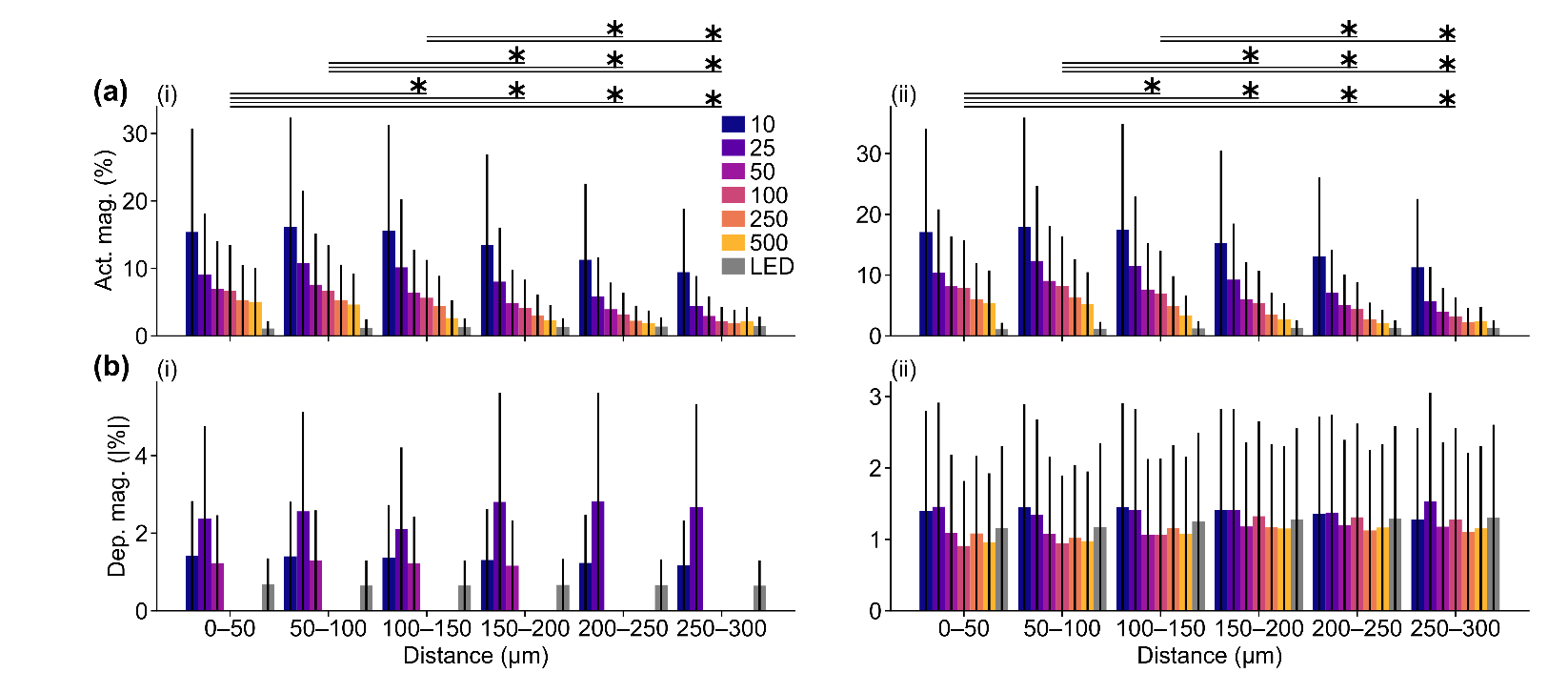

**Figure S7. Distance dependency of the mesoscopic-scale Ca^2+^ responses.**

Distance profiles of Ca^2+^ (a) activation and (b) depression (i) before and (ii) after the hemodynamic correction. Activation significantly different across distances (uncorrected Ca^2+^, two-way ANOVA, *F* = 43.78, *p* < 0.05; corrected Ca^2+^, *F* = 35.88, *p* < 0.05) and stimulation conditions (uncorrected Ca^2+^, *F* = 57.74, *p* < 0.05; uncorrected Ca^2+^, *F* = 57.27, *p* < 0.05). Depression was modulated by stimulation conditions (uncorrected Ca^2+^, *F* = 9.97, *p* < 0.05; corrected Ca^2+^, *F* = 4.78, *p* < 0.05) but stable across distances (uncorrected Ca^2+^, *F* = 0.18, *p* = 0.97; corrected Ca^2+^, *F* = 0.16, *p* = 0.98). Only activation exhibited significant interaction between distances and stimulation condition (uncorrected Ca^2+^, *F* = 2.00, *p* < 0.05; corrected Ca^2+^, *F* = 1.65, *p* < 0.05). * with horizontal line indicates *p* < 0.05 for pairwise comparison between distances with post-hoc Tukey’s HSD test.

**Table S1. Summary of post-hoc Tukey’s HSD test in Figure S5d**

| Condition 1 | Condition 2 | Difference of mean (%/s) | Adjusted p-value |
| --- | --- | --- | --- |
| 10 Hz | 25 Hz | -4.0 | < 0.05 |
| 10 Hz | 50 Hz | -18.1 | < 0.05 |
| 10 Hz | 100 Hz | -21.3 | < 0.05 |
| 10 Hz | 250 Hz | -20.8 | < 0.05 |
| 10 Hz | 500 Hz | -22.2 | < 0.05 |
| 10 Hz | LED | -21.1 | < 0.05 |
| 25 Hz | 50 Hz | -14.1 | < 0.05 |
| 25 Hz | 100 Hz | -17.3 | < 0.05 |
| 25 Hz | 250 Hz | -16.8 | < 0.05 |
| 25 Hz | 500 Hz | -18.2 | < 0.05 |
| 25 Hz | LED | -17.1 | < 0.05 |
| 50 Hz | 100 Hz | -3.2 | < 0.05 |
| 50 Hz | 250 Hz | -2.7 | < 0.05 |
| 50 Hz | 500 Hz | -4.1 | < 0.05 |
| 50 Hz | LED | -3.0 | < 0.05 |
| 100 Hz | 250 Hz | 0.5 | 0.99 |
| 100 Hz | 500 Hz | -0.9 | 0.78 |
| 100 Hz | LED | 0.2 | 1.00 |
| 250 Hz | 500 Hz | -1.4 | 0.32 |
| 250 Hz | LED | -0.3 | 1.00 |
| 500 Hz | LED | 1.1 | 0.75 |

**Table S2. Summary of post-hoc Tukey’s HSD test in Figure 9f across timings**

| Timing 1 | Timing 2 | Difference of mean | Adjusted p-value |
| --- | --- | --- | --- |
| Pre | Stim | -0.059 | < 0.05 |
| Pre | Post1 | -0.039 | < 0.05 |
| Pre | Post2 | -0.028 | < 0.05 |
| Pre | Post3 | -0.016 | 0.16 |
| Pre | Post4 | -0.006 | 0.97 |
| Pre | Post5 | 0.003 | 1.00 |
| Pre | Post6 | 0.011 | 0.62 |
| Stim | Post1 | 0.020 | < 0.05 |
| Stim | Post2 | 0.031 | < 0.05 |
| Stim | Post3 | 0.043 | < 0.05 |
| Stim | Post4 | 0.052 | < 0.05 |
| Stim | Post5 | 0.061 | < 0.05 |
| Stim | Post6 | 0.069 | < 0.05 |
| Post1 | Post2 | 0.011 | 0.57 |
| Post1 | Post3 | 0.023 | < 0.05 |
| Post1 | Post4 | 0.033 | < 0.05 |
| Post1 | Post5 | 0.042 | < 0.05 |
| Post1 | Post6 | 0.050 | < 0.05 |
| Post2 | Post3 | 0.012 | 0.45 |
| Post2 | Post4 | 0.022 | < 0.05 |
| Post2 | Post5 | 0.030 | < 0.05 |
| Post2 | Post6 | 0.039 | < 0.05 |
| Post3 | Post4 | 0.009 | 0.78 |
| Post3 | Post5 | 0.018 | 0.05 |
| Post3 | Post6 | 0.026 | < 0.05 |
| Post4 | Post5 | 0.009 | 0.82 |
| Post4 | Post6 | 0.017 | 0.09 |
| Post5 | Post6 | 0.008 | 0.88 |

**Table S3. Summary of post-hoc Tukey’s HSD test in Figure 9h across stimulation conditions**

| Condition 1 | Condition 2 | Difference of mean | Adjusted p-value |
| --- | --- | --- | --- |
| 10 Hz | 25 Hz | -0.015 | 0.16 |
| 10 Hz | 50 Hz | -0.011 | 0.50 |
| 10 Hz | 100 Hz | -0.001 | 1.00 |
| 10 Hz | 250 Hz | 0.017 | 0.09 |
| 10 Hz | 500 Hz | 0.019 | < 0.05 |
| 10 Hz | LED | 0.040 | < 0.05 |
| 25 Hz | 50 Hz | 0.004 | 1.00 |
| 25 Hz | 100 Hz | 0.015 | 0.21 |
| 25 Hz | 250 Hz | 0.032 | < 0.05 |
| 25 Hz | 500 Hz | 0.035 | < 0.05 |
| 25 Hz | LED | 0.055 | < 0.05 |
| 50 Hz | 100 Hz | 0.011 | 0.58 |
| 50 Hz | 250 Hz | 0.028 | < 0.05 |
| 50 Hz | 500 Hz | 0.031 | < 0.05 |
| 50 Hz | LED | 0.051 | < 0.05 |
| 100 Hz | 250 Hz | 0.018 | 0.06 |
| 100 Hz | 500 Hz | 0.020 | < 0.05 |
| 100 Hz | LED | 0.040 | < 0.05 |
| 250 Hz | 500 Hz | 0.003 | 1.00 |
| 250 Hz | LED | 0.023 | < 0.05 |
| 500 Hz | LED | 0.020 | 0.06 |

**Table S4. Summary of post-hoc Tukey’s HSD test in Figure 9h across distances**

| Distance 1 | Distance 2 | Difference of mean | Adjusted p-value |
| --- | --- | --- | --- |
| D1 | D2 | 0.002 | 1.00 |
| D1 | D3 | 0.013 | 0.37 |
| D1 | D4 | 0.024 | < 0.05 |
| D1 | D5 | 0.033 | < 0.05 |
| D1 | D6 | 0.043 | < 0.05 |
| D1 | D7 | 0.052 | < 0.05 |
| D2 | D3 | 0.011 | 0.54 |
| D2 | D4 | 0.023 | < 0.05 |
| D2 | D5 | 0.031 | < 0.05 |
| D2 | D6 | 0.041 | < 0.05 |
| D2 | D7 | 0.050 | < 0.05 |
| D3 | D4 | 0.012 | 0.47 |
| D3 | D5 | 0.020 | < 0.05 |
| D3 | D6 | 0.030 | < 0.05 |
| D3 | D7 | 0.039 | < 0.05 |
| D4 | D5 | 0.009 | 0.76 |
| D4 | D6 | 0.019 | < 0.05 |
| D4 | D7 | 0.028 | < 0.05 |
| D5 | D6 | 0.010 | 0.68 |
| D5 | D7 | 0.019 | < 0.05 |
| D6 | D7 | 0.009 | 0.75 |

**Table S5. Summary of post-hoc Tukey’s HSD test in Figure 11b**

| Condition 1 | Condition 2 | Difference of mean (%) | Adjusted p-value |
| --- | --- | --- | --- |
| 10 Hz | 25 Hz | 61.63 | < 0.05 |
| 10 Hz | 50 Hz | 8.58 | 0.73 |
| 10 Hz | 100 Hz | -20.86 | < 0.05 |
| 10 Hz | 250 Hz | -59.04 | < 0.05 |
| 10 Hz | 500 Hz | -84.28 | < 0.05 |
| 10 Hz | LED | -170.29 | < 0.05 |
| 25 Hz | 50 Hz | -53.05 | < 0.05 |
| 25 Hz | 100 Hz | -82.49 | < 0.05 |
| 25 Hz | 250 Hz | -120.66 | < 0.05 |
| 25 Hz | 500 Hz | -145.91 | < 0.05 |
| 25 Hz | LED | -231.91 | < 0.05 |
| 50 Hz | 100 Hz | -29.44 | < 0.05 |
| 50 Hz | 250 Hz | -67.62 | < 0.05 |
| 50 Hz | 500 Hz | -92.86 | < 0.05 |
| 50 Hz | LED | -178.87 | < 0.05 |
| 100 Hz | 250 Hz | -38.18 | < 0.05 |
| 100 Hz | 500 Hz | -63.42 | < 0.05 |
| 100 Hz | LED | -149.43 | < 0.05 |
| 250 Hz | 500 Hz | -25.25 | < 0.05 |
| 250 Hz | LED | -111.25 | < 0.05 |
| 500 Hz | LED | -86.00 | < 0.05 |

**Table S6. Summary of post-hoc Tukey’s HSD test in Figure 11d**

| Condition 1 | Condition 2 | Difference of mean (sec) | Adjusted p-value |
| --- | --- | --- | --- |
| 10 Hz | 25 Hz | 0.80 | < 0.05 |
| 10 Hz | 50 Hz | 0.35 | 0.54 |
| 10 Hz | 100 Hz | 0.40 | 0.38 |
| 10 Hz | 250 Hz | -0.55 | 0.07 |
| 10 Hz | 500 Hz | -2.28 | < 0.05 |
| 10 Hz | LED | -11.83 | < 0.05 |
| 25 Hz | 50 Hz | -0.45 | 0.24 |
| 25 Hz | 100 Hz | -0.40 | 0.38 |
| 25 Hz | 250 Hz | -1.36 | < 0.05 |
| 25 Hz | 500 Hz | -3.08 | < 0.05 |
| 25 Hz | LED | -12.63 | < 0.05 |
| 50 Hz | 100 Hz | 0.05 | 1.00 |
| 50 Hz | 250 Hz | -0.91 | < 0.05 |
| 50 Hz | 500 Hz | -2.63 | < 0.05 |
| 50 Hz | LED | -12.18 | < 0.05 |
| 100 Hz | 250 Hz | -0.96 | < 0.05 |
| 100 Hz | 500 Hz | -2.68 | < 0.05 |
| 100 Hz | LED | -12.23 | < 0.05 |
| 250 Hz | 500 Hz | -1.72 | < 0.05 |
| 250 Hz | LED | -11.27 | < 0.05 |
| 500 Hz | LED | -9.55 | < 0.05 |

**Table S7. Summary of post-hoc Tukey’s HSD test in Figure 11f**

| Condition 1 | Condition 2 | Difference of mean (%) | Adjusted p-value |
| --- | --- | --- | --- |
| 10 Hz | 25 Hz | 1.03 | 0.08 |
| 10 Hz | 50 Hz | 0.28 | 0.99 |
| 10 Hz | 100 Hz | -0.77 | 0.36 |
| 10 Hz | 250 Hz | -2.05 | < 0.05 |
| 10 Hz | 500 Hz | -2.75 | < 0.05 |
| 10 Hz | LED | -3.40 | < 0.05 |
| 25 Hz | 50 Hz | -0.76 | 0.39 |
| 25 Hz | 100 Hz | -1.80 | < 0.05 |
| 25 Hz | 250 Hz | -3.08 | < 0.05 |
| 25 Hz | 500 Hz | -3.78 | < 0.05 |
| 25 Hz | LED | -4.43 | < 0.05 |
| 50 Hz | 100 Hz | -1.05 | 0.07 |
| 50 Hz | 250 Hz | -2.33 | < 0.05 |
| 50 Hz | 500 Hz | -3.02 | < 0.05 |
| 50 Hz | LED | -3.67 | < 0.05 |
| 100 Hz | 250 Hz | -1.28 | < 0.05 |
| 100 Hz | 500 Hz | -1.98 | < 0.05 |
| 100 Hz | LED | -2.63 | < 0.05 |
| 250 Hz | 500 Hz | -0.69 | 0.50 |
| 250 Hz | LED | -1.34 | < 0.05 |
| 500 Hz | LED | -0.65 | 0.73 |

**Table S8. Summary of post-hoc Tukey’s HSD test in Figure 11h**

| Condition 1 | Condition 2 | Difference of mean (sec) | Adjusted p-value |
| --- | --- | --- | --- |
| 10 Hz | 25 Hz | 3.33 | < 0.05 |
| 10 Hz | 50 Hz | 2.34 | < 0.05 |
| 10 Hz | 100 Hz | 0.80 | 0.48 |
| 10 Hz | 250 Hz | -0.73 | 0.61 |
| 10 Hz | 500 Hz | -1.83 | < 0.05 |
| 10 Hz | LED | -2.92 | < 0.05 |
| 25 Hz | 50 Hz | -1.00 | 0.22 |
| 25 Hz | 100 Hz | -2.53 | < 0.05 |
| 25 Hz | 250 Hz | -4.06 | < 0.05 |
| 25 Hz | 500 Hz | -5.16 | < 0.05 |
| 25 Hz | LED | -6.25 | < 0.05 |
| 50 Hz | 100 Hz | -1.53 | < 0.05 |
| 50 Hz | 250 Hz | -3.06 | < 0.05 |
| 50 Hz | 500 Hz | -4.16 | < 0.05 |
| 50 Hz | LED | -5.26 | < 0.05 |
| 100 Hz | 250 Hz | -1.53 | < 0.05 |
| 100 Hz | 500 Hz | -2.63 | < 0.05 |
| 100 Hz | LED | -3.72 | < 0.05 |
| 250 Hz | 500 Hz | -1.10 | 0.13 |
| 250 Hz | LED | -2.19 | < 0.05 |
| 500 Hz | LED | -1.09 | 0.28 |

**Table S9. Summary of post-hoc Tukey’s HSD test in Figure 11j**

| Condition 1 | Condition 2 | Difference of mean (a.u.) | Adjusted p-value |
| --- | --- | --- | --- |
| 10 Hz | 25 Hz | 0.31 | 0.46 |
| 10 Hz | 50 Hz | 0.88 | < 0.05 |
| 10 Hz | 100 Hz | 1.17 | < 0.05 |
| 10 Hz | 250 Hz | 1.41 | < 0.05 |
| 10 Hz | 500 Hz | 1.67 | < 0.05 |
| 10 Hz | LED | 0.31 | 0.63 |
| 25 Hz | 50 Hz | 0.57 | < 0.05 |
| 25 Hz | 100 Hz | 0.85 | < 0.05 |
| 25 Hz | 250 Hz | 1.09 | < 0.05 |
| 25 Hz | 500 Hz | 1.36 | < 0.05 |
| 25 Hz | LED | 0.00 | 1.00 |
| 50 Hz | 100 Hz | 0.29 | 0.56 |
| 50 Hz | 250 Hz | 0.53 | < 0.05 |
| 50 Hz | 500 Hz | 0.79 | < 0.05 |
| 50 Hz | LED | -0.57 | < 0.05 |
| 100 Hz | 250 Hz | 0.24 | 0.76 |
| 100 Hz | 500 Hz | 0.50 | < 0.05 |
| 100 Hz | LED | 0.85 | < 0.05 |
| 250 Hz | 500 Hz | 0.26 | 0.66 |
| 250 Hz | LED | -1.09 | < 0.05 |
| 500 Hz | LED | -1.36 | < 0.05 |

**Table S10. Summary of post-hoc Tukey’s HSD test in Figure 11l**

| Condition 1 | Condition 2 | Difference of mean (a.u.) | Adjusted p-value |
| --- | --- | --- | --- |
| 10 Hz | 25 Hz | 4.06 | < 0.05 |
| 10 Hz | 50 Hz | -0.67 | 0.95 |
| 10 Hz | 100 Hz | -4.09 | < 0.05 |
| 10 Hz | 250 Hz | -7.91 | < 0.05 |
| 10 Hz | 500 Hz | -9.57 | < 0.05 |
| 10 Hz | LED | -17.59 | < 0.05 |
| 25 Hz | 50 Hz | -4.73 | < 0.05 |
| 25 Hz | 100 Hz | -8.15 | < 0.05 |
| 25 Hz | 250 Hz | -11.97 | < 0.05 |
| 25 Hz | 500 Hz | -13.63 | < 0.05 |
| 25 Hz | LED | -21.65 | < 0.05 |
| 50 Hz | 100 Hz | -3.41 | < 0.05 |
| 50 Hz | 250 Hz | -7.23 | < 0.05 |
| 50 Hz | 500 Hz | -8.90 | < 0.05 |
| 50 Hz | LED | -16.91 | < 0.05 |
| 100 Hz | 250 Hz | -3.82 | < 0.05 |
| 100 Hz | 500 Hz | -5.48 | < 0.05 |
| 100 Hz | LED | -13.50 | < 0.05 |
| 250 Hz | 500 Hz | -1.67 | 0.31 |
| 250 Hz | LED | -9.68 | < 0.05 |
| 500 Hz | LED | -8.02 | < 0.05 |

**Supplementary movies**

Movie S1: Time-lapse mesoscopic-scale images of Ca^2+^ signal in the 10-Hz ICMS condition (same mouse as Figure 2).

Movie S2: Time-lapse mesoscopic-scale images of Ca^2+^ signal in the 100-Hz ICMS condition (same mouse as Figure S2).

Movie S3: Time-lapse mesoscopic-scale images of Ca^2+^ signal in the visual stimulation condition (same mouse as Figure 3).

Movie S4: Time-lapse mesoscopic-scale images of oxyhemoglobin signal in the 10-Hz ICMS condition (same mouse as Figure 9).

Movie S5: Time-lapse mesoscopic-scale images of oxyhemoglobin signal in the 100-Hz ICMS condition (same mouse as Figure 9).

Movie S6: Time-lapse mesoscopic-scale images of oxyhemoglobin signal in the visual stimulation condition (same mouse as Figure 9).

Movie S7: Time-lapse mesoscopic-scale images of deoxyhemoglobin signal in the 10-Hz ICMS condition (same mouse as Figure 9).

Movie S8: Time-lapse mesoscopic-scale images of deoxyhemoglobin signal in the 100-Hz ICMS condition (same mouse as Figure 9).

Movie S9: Time-lapse mesoscopic-scale images of deoxyhemoglobin signal in the visual stimulation condition (same mouse as Figure 9).

Movie S10: Time-lapse two-photon images of Ca^2+^ signal in the 10-Hz ICMS condition (same mouse as Figure 11a–e).

Movie S11: Time-lapse two-photon images of Ca^2+^ signal in the 100-Hz ICMS condition (same mouse as Figure 11a–e).

Movie S12: Time-lapse two-photon images of Ca^2+^ signal in the visual stimulation condition (same mouse as Figure 11a–e).

Stimulation train was applied from t = 0 to 10 s (annotation at upper left corner).
